# Exploring the antimicrobial potential of streptococcal fratricins

**DOI:** 10.64898/2026.10.23.761373

**Authors:** Rebekka Moe, Morten Kjos, Daniel Straume

**Affiliations:** Faculty of Chemistry, Biotechnology and Food Science, Norwegian University of Life Sciences, Ås, Norway

## Abstract

Many streptococci are competent for natural transformation, during which they express a murein hydrolase that lyses and kills closely related species, thereby making homologous DNA available for uptake by competent cells. This process, termed fratricide, is mediated by enzymes known as fratricins. The most extensively studied fratricin, CbpD from *Streptococcus pneumoniae*, is highly specific, efficient, and rapid in its killing, highlighting the potential of fratricins as targeted antimicrobial agents. Here, we screened nine putative fratricins to evaluate their antimicrobial activity. Of these, Zoocin A from *Streptococcus agalactiae*, ScrM from *Streptococcus dysgalactiae*, and CbpD from *Streptococcus vestibularis* were successfully purified. Zoocin A exhibited the broadest target range, whereas ScrM and CbpD-ves showed somewhat narrower activity. All three enzymes predominantly targeted streptococci. Notably, Zoocin A was active against *Streptococcus mutans* and *Streptococcus sobrinus*, species associated with dental caries. The enzymes were also active against *S. agalactiae*, *S. dysgalactiae*, and *S. pyogenes*, which are associated with severe infections in humans and animals. We characterised the activity and stability of Zoocin A and ScrM and found that both retained activity under physiologically relevant conditions of pH, temperature, and ionic strength. ScrM activity was independent of divalent cations, whereas Zoocin A activity required cations and was observed with both Ca^2+^ and Mn^2+^. For both enzymes, peptidoglycan cleavage was localised to the septal region of dividing cells. Zoocin A shows promise as an antimicrobial agent against *S. mutans*, but its efficacy against biofilms would likely benefit from further optimisation or combination with biofilm-disrupting agents.

**Importance:** Streptococci are major pathogens of humans and animals, causing infections ranging from common mucosal and dental diseases to severe invasive and systemic infections. The increasing prevalence of antimicrobial resistance among clinically important streptococci is a major challenge to healthcare systems who are running out of treatment options. It is therefore necessary to develop new approaches to prevent and treat streptococcal infections. Fratricins are streptococcal peptidoglycan hydrolases that mediate killing of closely related bacteria during natural competence, suggesting that they could be harnessed as targeted antimicrobials against pathogenic streptococci. However, the diversity and antimicrobial potential of fratricins remain poorly explored. By identifying and characterising putative fratricins, this work contributes to expanding the repertoire of antimicrobials and provides a foundation for developing fratricin-based approaches to combat streptococcal infections.

## Introduction

The continued emergence and spread of antibiotic resistance highlight the urgent need to identify and develop new antimicrobial agents. Pathogenic streptococci remain major causes of infection in humans and animals worldwide. Although many streptococci remain susceptible to β-lactams, resistance to alternative antimicrobial agents, particularly macrolides and lincosamides, represents a substantial global concern, and strategies that prevent infections in the first place would reduce antibiotic use and the spread of resistance (Shivaee et al., 2025; World Health Organization, 2024). Furthermore, alternatives to β-lactams are particularly important for patients with β-lactam allergies and for treating biofilm-associated infections, where conventional antibiotic therapy is often less effective.

Streptococci are a diverse group of Gram-positive bacteria that commonly colonise mucosal surfaces and include numerous opportunistic and primary pathogens of both humans and animals. Based on phylogenetic relationships, streptococci are divided into several groups, including the mitis, anginosus, salivarius, sanguinis, bovis, and pyogenic groups, which differ in their ecological niches, host associations, and pathogenic potential (Kawamura et al., 1995; Richards et al., 2014). *Streptococcus pneumoniae*, an important human pathogen responsible for respiratory and invasive infections, belongs to the mitis group. Streptococci are competent for natural transformation, meaning that under certain conditions they express genes that enable them to scavenge for, take up, and incorporate naked DNA into their genomes, thereby acquiring new genetic traits. Competence is thought to be a bacterial stress response that enhances adaptability under hostile conditions (Engelmoer & Rozen, 2011; Maziero et al., 2023; Prudhomme et al., 2024). For example, antibiotic exposure can trigger the competent state, which may allow the bacteria to acquire resistance genes (Prudhomme et al., 2024). Natural transformation was first discovered in *S. pneumoniae* almost 100 years ago and has since been described in both Gram-positive and Gram-negative bacteria across a wide range of phyla (Johnston et al., 2014). In streptococci, the competent state is transient and defined as a two-part process: the first (early) stage is the induction of genes required to enter the competent state, and the second (late) stage is the expression of genes involved in transformation itself (Luo & Morrison, 2003; Peterson et al., 2000). Streptococci possess either of two systems for competence induction: the ComCDE or ComRS system (Fontaine et al., 2010; Håvarstein et al., 1996; Pestova et al., 1996). Although the two systems differ in several respects, they both ultimately result in the production of the alternative sigma factor ComX (SigX in some species). Expression of ComX marks the entry into the late competent stage, in which it induces transcription of so-called late competence genes, including genes involved in DNA uptake and recombination (Peterson et al., 2000; Peterson et al., 2004). Several streptococcal species encode a peptidoglycan hydrolase with a ComX-inducible promoter (Berg, Ohnstad, et al., 2012). CbpD from *S. pneumoniae* is the most studied among these and has been shown to increase transformation rates by lysing closely related species, thereby making homologous DNA available to the recipient cell (Johnsborg et al., 2008; Kausmally et al., 2005). This process has been termed *fratricide*, and the acting hydrolase a *fratricin*. To avoid suicide, competent pneumococci express an immunity protein, ComM, that protects the cells against lysis (Håvarstein et al., 2006; Juillot et al., 2025; Minhas et al., 2023; Straume et al., 2017).

There are two main types of these putative fratricins, CbpD-like or LytF, and most streptococci encode either one of these enzymes (Berg, Biørnstad, et al., 2012; Berg, Ohnstad, et al., 2012). Common for all, is that they contain a cell wall hydrolytic domain in addition to domains required for binding to the surface of susceptible target cells. The variability among cell-surface-binding domains in these enzymes likely reflects differences in target cell recognition. The domain composition of the CbpD-like hydrolases varies between species, but all contain an N-terminal hydrolytic cysteine/histidine-dependent amidohydrolase/peptidase (CHAP) domain, which is usually followed by either one or two Src homology 3b (SH3b) domains. The C-terminal domains of CbpDs show more diversity. CbpDs encoded by members of the mitis phylogenetic group contain a C-terminal choline-binding domain (Cbd), which is where the name of the enzyme originates from, while some members of the pyogenic group and all members of the salivarius group contain a C-terminal conserved domain (CCD) in addition to or instead of the SH3b domains, respectively. Recently, CCD was shown to be essential for target cell binding of the putative fratricin ScrM expressed by *Streptococcus dysgalactiae* (Mårli et al., 2025). ScrM and its cognate immunity protein ScrI have also been proposed to constitute a fratricin-immunity factor system analogous to CbpD-ComM in *S. pneumoniae* (Mårli et al., 2025). However, its involvement in fratricide has not yet been demonstrated. The LytF (putative) fratricins contain a C-terminal CHAP domain and three to five group B streptococcal secreted protein (Bsp) domains in the N-terminal part. Evidence suggests that LytF in *Streptococcus mutans* and *Streptococcus gordonii* functions as a fratricin analogous to CbpD from *S. pneumoniae* (Berg, Ohnstad, et al., 2012; Eaton & Jacques, 2010), although this appears not to be the case in *Streptococcus sanguinis,* in which it instead has been shown to increase the transformation rate by facilitating extrusion of the transformation pilus (Moe et al., 2026). The CbpD-like fratricins in *Streptococcus thermophilus* (CbpD-St) and *Streptococcus suis* (CrfP) have also been shown to increase the transformation rate of the producer, but how they do this has not yet been demonstrated (Bjørnstad et al., 2012; Zhu et al., 2021). An overview of fratricins found across the streptococcal phylogenetic tree is shown in Fig. 1.

**Fig. 1.**
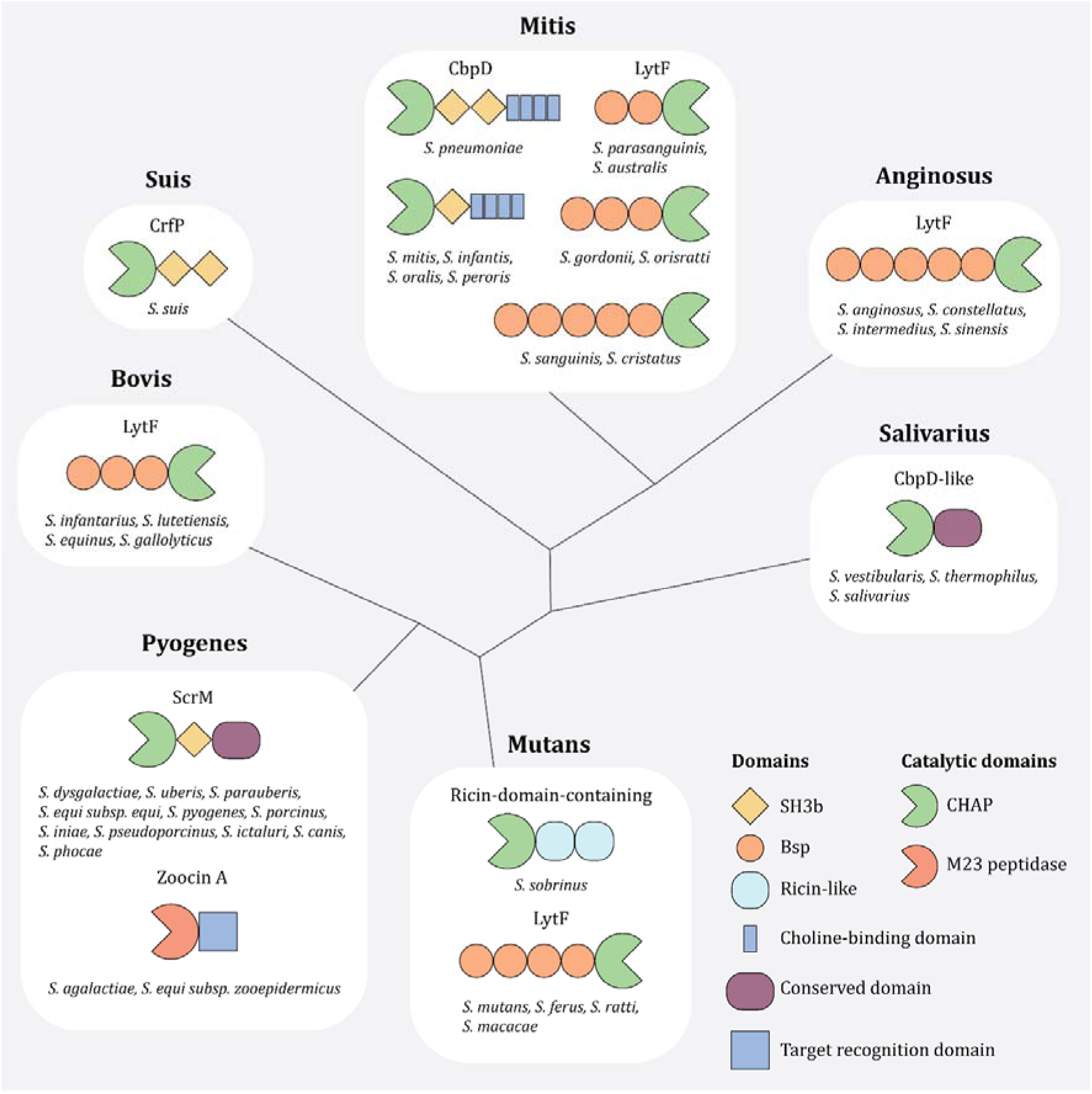
Domain organisation of putative fratricins across streptococcal phylogenetic groups. Putative fratricins were identified based on homology to known CHAP-domain sequences in BLASTp searches or on previous identification (Berg, Ohnstad, et al., 2012). Protein domains were predicted using InterProScan 6 (Blum et al., 2026). The connecting lines illustrate phylogenetic relationships but do not represent evolutionary distances. The amino acid sequences used for structural prediction are provided in Supplemental material S2. Abbreviations: SH3b, Src homology 3b; Bsp, group B streptococcal secreted protein; CHAP, cysteine/histidine-dependent amidohydrolase/peptidase.

Streptococcal fratricins constitute a somewhat understudied group of peptidoglycan hydrolases, and detailed functional characterisation is currently available only for a limited number of representatives. Among these, CbpD from *S. pneumoniae* (CbpD-Sp) is the most studied and therefore provides an important model for the biology of fratricins. If other fratricins exhibit similar properties, they may hold promise as antimicrobial agents. CbpD-Sp displays a narrow target spectrum, acting only against its immediate relatives with choline-covered teichoic acids, *Streptococcus mitis* and *Streptococcus oralis* (Johnsborg et al., 2008). This specificity likely reflects the ecological role of fratricins in mediating the release of homologous DNA. It has been shown for the CbpD homologue in *S. mitis* B6 that addition of purified enzyme to cell cultures resulted in rapid and efficient lysis (Straume et al., 2020), illustrating the potential of fratricins as highly precise antimicrobial enzymes.

The aim of this study was to expand the repertoire of antimicrobials by investigating streptococcal fratricins as a potential new class of enzyme-based antimicrobials targeting pathogenic streptococci. The enzymes investigated in this study were selected based on their production by streptococcal species outside the Pneumoniae subclade of the mitis phylogenetic group (Patel & Gupta, 2018) as they may represent promising candidates for activity against non-pneumococcal streptococci. *S. pneumoniae* is an important cause of respiratory and invasive infections, whereas many non-pneumococcal streptococci are associated with colonisation and infection at mucosal and other exposed sites (Lannes Costa et al., 2021; Narciso et al., 2025). While antimicrobial enzymes have potential as therapeutic agents, their use for systemic treatment is limited by factors such as proteolytic degradation and interactions with host components. Their application may be better suited in localised or surface-associated settings. For example, antimicrobial enzymes could be used to coat plastic or metal surfaces to prevent bacterial adhesion and biofilm formation, thereby reducing the risk of device-associated infections. This approach could apply to both medical and veterinary settings, including the development of antimicrobial surfaces for indwelling devices and other contact materials.

We screened nine putative fratricins across all streptococcal phylogenetic groups to assess their potential as antimicrobial agents. Of these, Zoocin A from *Streptococcus agalactiae*, ScrM from *S. dysgalactiae*, and CbpD from *Streptococcus vestibularis* were successfully purified. We then characterised Zoocin A and ScrM for potential antimicrobial use, including activity against species within and outside the *Streptococcus* genus, stability and activity under different conditions, and potential for biofilm-related applications.

## Results

### Screening putative fratricins for antimicrobial potential

Based on the properties of CbpD-sp (and the *S. mitis* homologue), fratricins may represent a group of enzymes with great potential as antimicrobial agents against pathogenic streptococci. Following an analysis of predicted domain architectures of putative fratricins (Fig. 1), we selected nine putative fratricins from the following nine species: *S. sanguinis*, *S. mutans*, *S. sobrinus*. *S. anginosus*, *S. dysgalactiae*, *S. agalactiae*, *S. vestibularis*, *S. suis*, and *S. equinus*. This selection aimed to cover all major streptococcal phylogenetic groups and to include variety in the predicted structure and target range of the proteins. *S. dysgalactiae*, *S. vestibularis*, and *S. suis* encode CbpD-like proteins, and *S. sanguinis*, *S. mutans*, *S. anginosus*, and *S. equinus* encode LytF proteins. *S. agalactiae* encodes Zoocin A, a smaller and unrelated enzyme consisting of an M23 peptidase domain and a target recognition domain (TRD). *S. sobrinus* encodes a unique putative fratricin, predicted to consist of an M23 peptidase domain and two Ricin-like domains. The corresponding genes were fused N-terminally with a His-tag, expressed in *Escherichia coli* and purified using immobilised metal ion affinity chromatography. We were unable to construct plasmids carrying the *lytF* genes from *S. sanguinis* and *S. equinus*. Truncated versions of the genes identified by sequencing suggest that the full-length *lytF* from these species are toxic to *E. coli*. ScrM (*S. dysgalactiae*) was not included in this screen because it has been successfully purified in our lab previously. We were able to purify significant amounts of the putative fratricins of *S. agalactiae* (Zoocin A), *S. vestibularis* (CbpD-Sv), and *S. sobrinus* (NCBI RefSeq WP_019775801.1) (Supplemental Fig. S1). On the other hand, purification of LytF proteins was not successful using this approach.

Zoocin A, ScrM, and CbpD-Sv were selected for assessment of their antimicrobial activity and target range by treating exponentially growing cells with purified enzymes (30 µg/mL). Enzymes were considered active against a given species when the OD_550_ decreased by more than 50% relative to the control. A decrease of 25-50% was classified as partial activity, whereas a decrease of less than 25% was classified as no detectable activity. The results are summarized in Table 1. For ScrM data from this study is combined with previously published results by Mårli et al. (2025). Zoocin A showed broad-spectrum activity against streptococcal species, killing 10 of the 14 species tested (Table 1). Species susceptible to Zoocin A were found in all phylogenetic groups, except the suis group. By comparison, ScrM exhibited a slightly narrower range of susceptible species, killing 6 of the 14 streptococcal species, which belong to the salivarius, pyogenes, and anginosus groups (Table 1). Similarly, CbpD-Sv was active against 7 out of 14 streptococcal species, belonging to the mutans, salivarius, bovis, and pyogenes groups (Table 1). Notably, *S. vestibularis* demonstrated complete resistance to CbpD-Sv, in contrast to *S. agalactiae* and *S. dysgalactiae*, which were highly susceptible to their respective putative fratricins. The three enzymes were also tested for activity against a selection of non-streptococcal pathogens. Both Zoocin A and ScrM were active against *Enterococcus faecalis*, but neither was active against any of the other non-streptococcal species.

**Table 1.** Target range of Zoocin A, ScrM, and CbpD-Sv. Activity classifications are based on data shown in Supplemental Fig. S2, together with previously published data by Mårli et al. (2025).

| Species<sup>1</sup> | Phylogenetic group (streptococci) | Zoocin A | ScrM | CbpD-Sv |
| --- | --- | --- | --- | --- |
| *S. mitis* | Mitis | No | No<sup>2</sup> | No |
| *S. pneumoniae* |  | No | No<sup>2</sup> | No |
| *S. sanguinis* |  | No | No | No |
| *S. cristatus* |  | Yes | No<sup>2</sup> | No |
| *S. mutans* | Mutans | Yes | No<sup>2</sup> | Partial |
| *S. sobrinus* |  | Yes | No<sup>2</sup> | Partial |
| *S. thermophilus* | Salivarius | Yes | Yes<sup>2</sup> | Yes |
| *S. vestibularis* |  | Yes | Yes<sup>2</sup> | No |
| *S. equinus* | Bovis | Yes | No<sup>2</sup> | Yes |
| *S. suis* | Suis | No | No<sup>2</sup> | No |
| *S. dysgalactiae* | Pyogenes | Yes | Yes<sup>2</sup> | Yes |
| *S. agalactiae* |  | Yes | Yes<sup>2</sup> | Yes |
| *S. pyogenes* |  | Yes | Yes<sup>2</sup> | Yes |
| *S. anginosus* | Anginosus | Partial<sup>3</sup> | Yes | No |
| *E. faecium* | - | No | No | No |
| *E. faecalis* | - | Partial | Partial | No |
| *L. innocua* | - | No | No | No |
| *L. garviae* | - | No | No | No |
| *S. haemolyticus* | - | No | No | No |
| *S. aureus* | - | No | No | No |
| *B. subtilis* | - | No | No | No |

<sup>1</sup> Strain information is provided in Supplemental Table S1.
<sup>2</sup> Mårli et al., 2025
<sup>3</sup> This species was classified as partially susceptible due to high variability between the replicates.

### Activity and stability profiling of Zoocin A and ScrM

Zoocin A and ScrM were selected for further characterisation because they have different catalytic and target recognition domains in addition to representing a broad- and narrow-spectrum enzyme, respectively. In all tests described in this section *S. agalactiae* was used as target cells for Zoocin A activity and *S. dysgalactiae* for ScrM.

To assess the heat tolerance of the enzymes, they were incubated at different temperatures for 1 h before being added to actively growing cultures at 37°C. Both enzymes retained full activity following pre-treatment at temperatures up to 37°C, resulting in nearly complete lysis of the cultures, whereas incubation at 45°C effectively abolished enzymatic activity (Fig. 2).

**Fig. 2.**
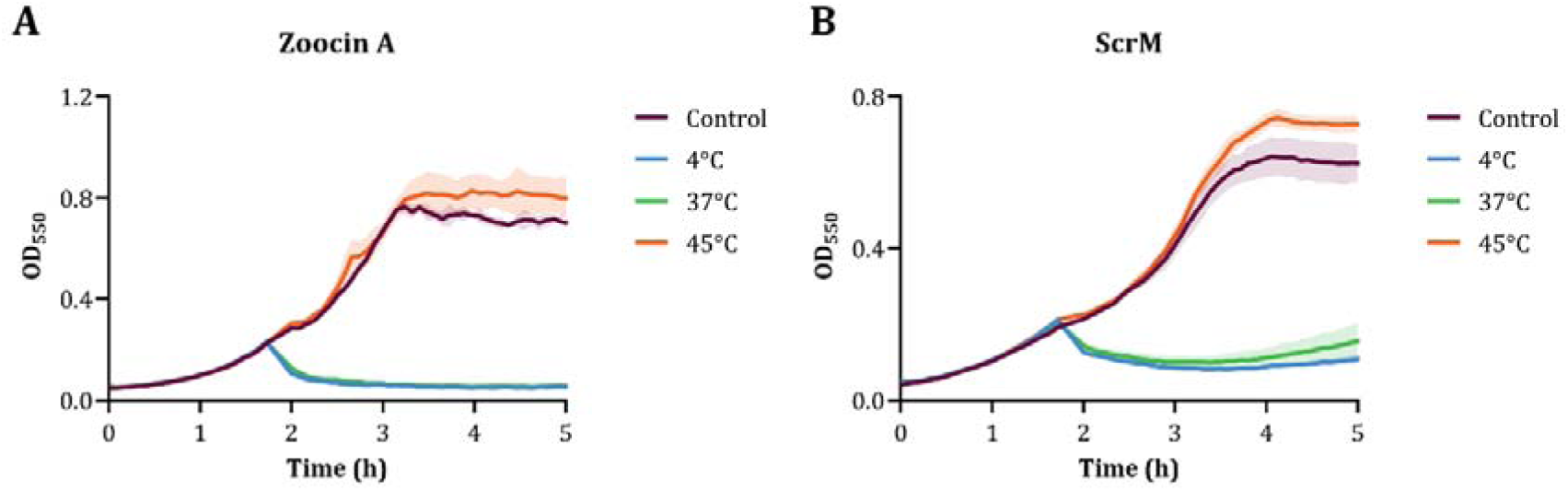
Temperature tolerance of Zoocin A (A) and ScrM (B). The enzymes were incubated for 1 hour at 4°C, 37°C, and 45°C before being added to cultures growing at 37°C at an OD_550_ of 0.2 to a final concentration of 2 µg/mL.

Next, to assess the enzymes’ ionic strength tolerance, we sought to perform the assays in a PBS or TBS buffer in which the salt (NaCl) concentration could be precisely controlled. However, lytic activity was reduced in these buffers compared with that observed in C-medium, independent of NaCl concentration (Supplemental Fig. S3). This effect was particularly pronounced for ScrM, which reduced culture turbidity only by approximately 20% after treatment at the MIC_50_ concentration (Supplemental Fig. S3; see Supplementary Fig. S4 for MIC_50_ determination). Therefore, assays to determine salt tolerance were conducted in C-medium supplemented with NaCl to the indicated concentrations (thus, the stated concentrations represent the added NaCl and do not include the salt inherently present in the growth medium). Zoocin A exhibited an almost linear loss of enzymatic activity with increasing NaCl levels but did not become fully inactive even at the highest concentration (1 M) (Fig. 3A). Mean culture turbidity increased from 22% in the unsupplemented medium to approximately 60% at 0.25 M NaCl and 90% at 1 M NaCl (Fig. 3A). Although ScrM activity decreased with increasing NaCl concentration, the effect was less pronounced than that observed for Zoocin A (Fig. 3B). Mean culture turbidity increased from 35% in the absence of added NaCl to 43% at 0.25 M NaCl and 78% at 1 M NaCl (Fig. 3B). Supplemental Fig. S5 and S6 show the corresponding growth curves for these experiments.

**Fig. 3.**
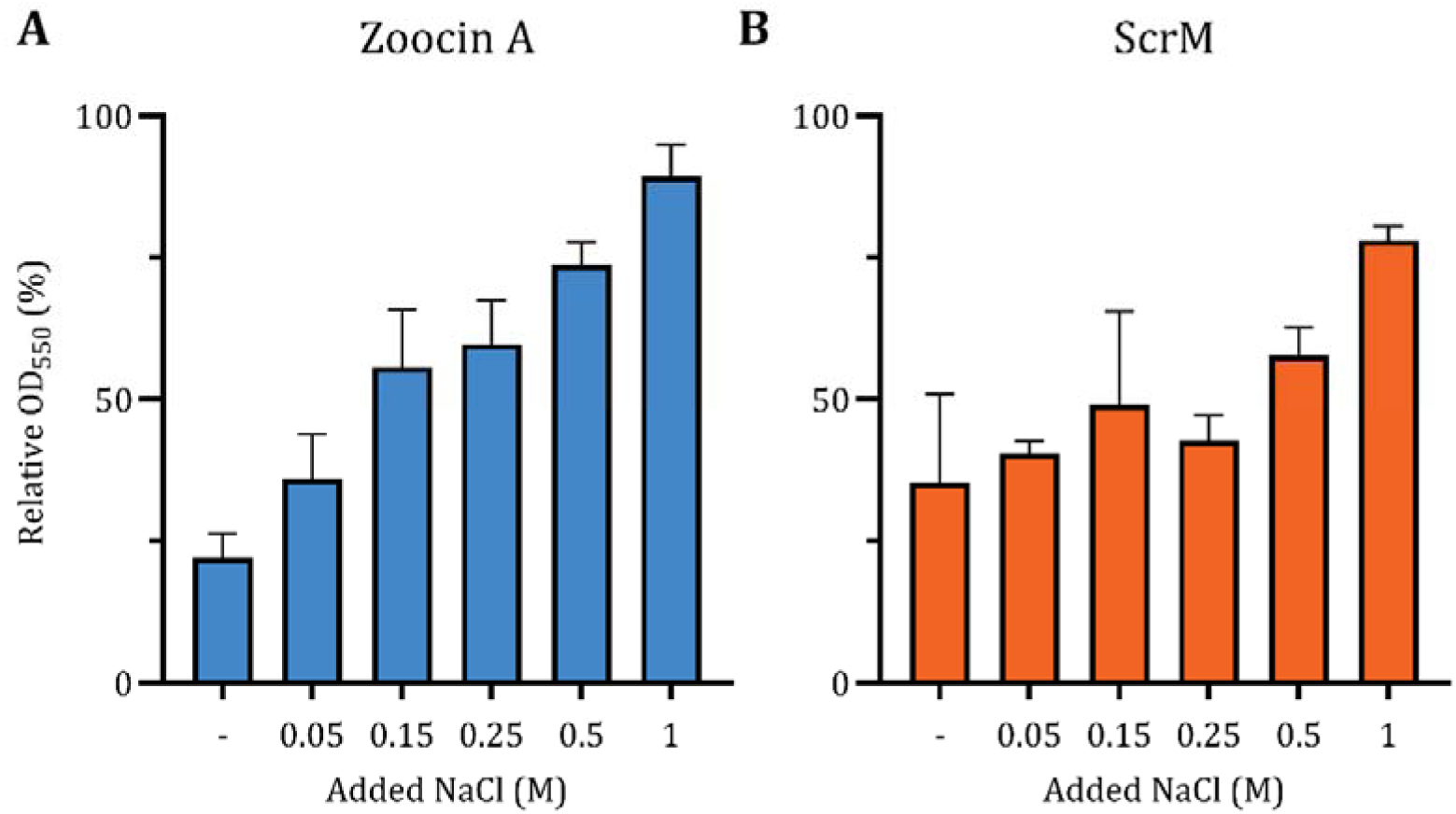
Enzymatic activity of Zoocin A (A) and ScrM (B) at increasing concentrations of NaCl. Cultures of *S. agalactiae* (Zoocin A) and *S. dysgalactiae* (ScrM) were supplemented with the indicated NaCl concentrations at OD_550_ ∼ 0.2. Enzyme activity was assessed after addition of enzymes pre-incubated in the corresponding NaCl concentrations. The enzymes were added to a final concentration of 4 µg/mL. Cultures with no added NaCl (-) were included as controls and were treated with enzymes incubated in dH_2_O. The relative OD_550_ was determined by comparing the OD_550_ measured 1 h after enzyme addition with the OD_550_ immediately before treatment.

The effect of pH on enzyme activity was subsequently investigated. To avoid potential differences in bacterial growth resulting from the experimental pH values, cultures were grown to an OD_550_ of 0.2 in standard C-medium, then harvested and resuspended in pH-adjusted media immediately before enzyme addition. pH values ranging from 3 to 10 were tested. Zoocin A retained activity at pH from 7 to 9, with the highest lytic effect at pH 8 (Fig. 4A). ScrM was active at pH ranging from 6 to 9, with the highest activity at pH 7 to 9 (Fig. 4B).

**Fig. 4.**
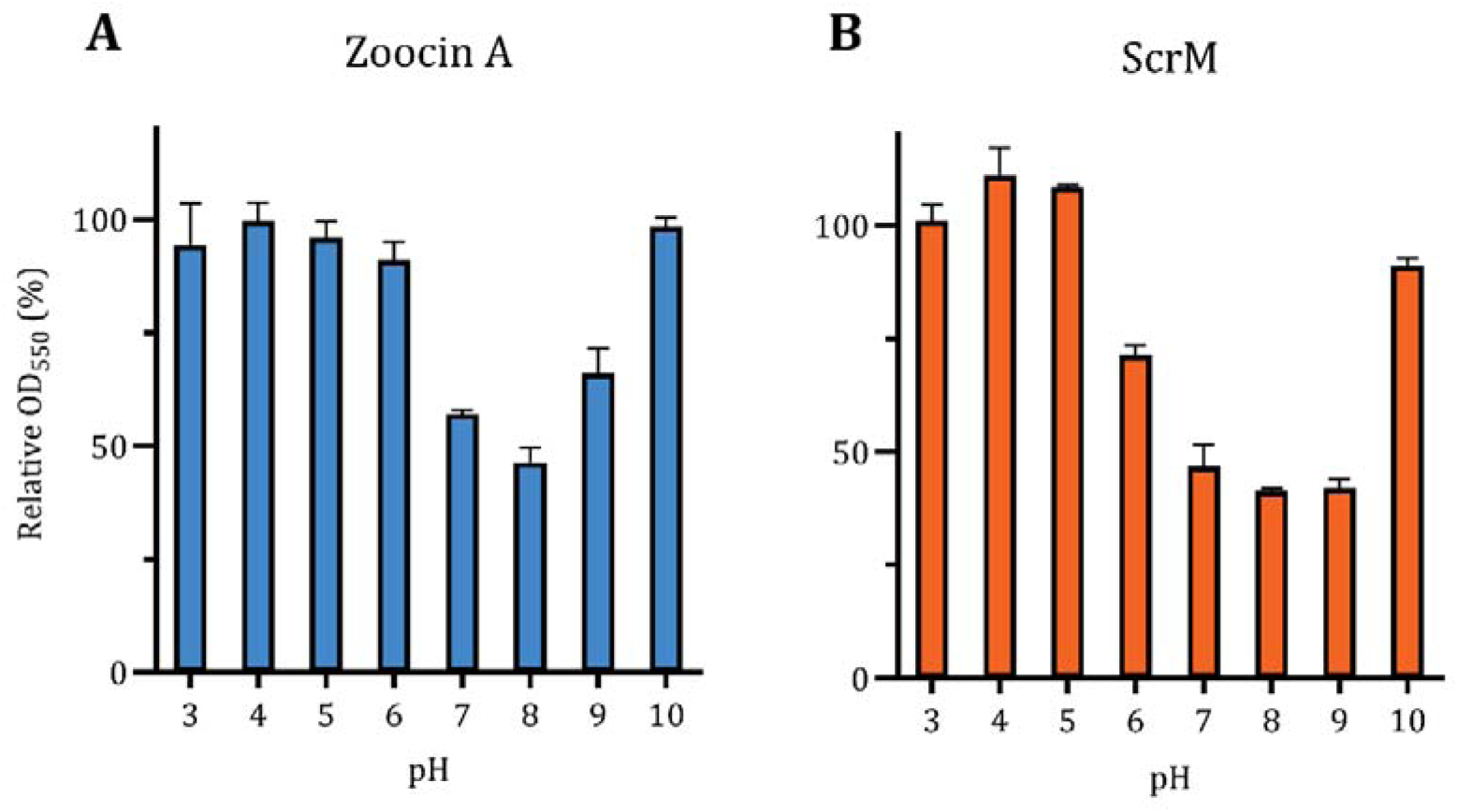
Enzymatic activity of Zoocin A (A) and ScrM (B) across pH ranging from 3 to10. Cells of *S. agalactiae* and *S. dysgalactiae* were grown to an OD_550_ of ∼ 0.2 before their medium was replaced with pH-adjusted C-medium. To avoid measuring lytic effects induced by non-physiological pH values, enzyme activity was determined by measuring the cultures’ decrease in OD_550_ 5 minutes after enzyme addition. A final concentration of 4 µg/mL enzyme were added.

Since it has previously been shown that the chelating agent EDTA increased the lytic effect of CbpD-sp (Eldholm et al., 2009), we tested how EDTA and various divalent cations influenced the lytic activity of Zoocin A and ScrM. Adding EDTA to the growth medium and EDTA pre-treatment of the enzymes increased the lytic effect of both enzymes when added to actively growing cells (Supplemental Fig. S7). This could suggest that neither enzyme requires divalent cations for its activity. However, the increased lytic effect may also reflect changes in cell susceptibility from the EDTA treatment or incomplete chelation of divalent cations in the growth medium. To investigate this further, we assessed the enzymes’ metal-ion dependence by zymography. Following EDTA treatment of the zymogram gels, Zoocin A did not produce clearing zones in the cell-embedded gel, but activity was restored upon addition of Mn^2+^ or Ca^2+^, whereas no activity was observed following addition of Zn^2+^ (Fig. 5A). This suggests that Zoocin A requires divalent cations for its activity and can utilise both Mn^2+^ and Ca^2+^. In contrast, ScrM formed clearing zones despite EDTA treatment, suggesting it does not require divalent cations for its enzymatic activity (Fig. 5B).

**Fig. 5.**
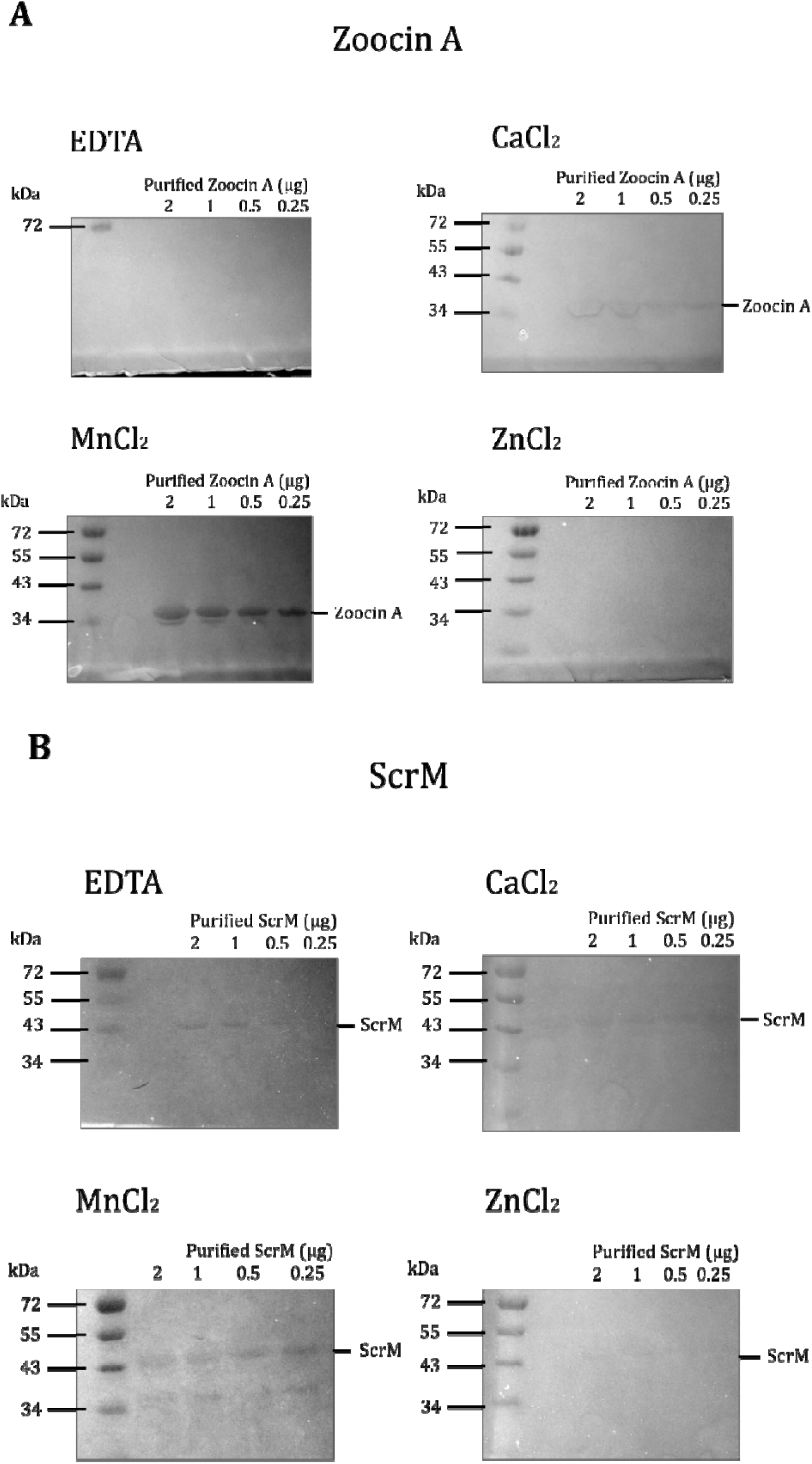
Effects of the divalent cations Ca²□, Mn²□, and Zn²□on the activity of Zoocin A (A) and ScrM (B) assessed by zymography. Cell-embedded SDS-PAGE gels containing Zoocin A or ScrM (2, 1, 0.5, and 0.25 µg) were treated with 1 mM EDTA to remove divalent cations and subsequently submerged in refolding buffers containing 1 mM CaC, MnCl, or ZnCl to assess the effect of each cation on enzyme activity. Gels maintained in EDTA were included as controls. For Zoocin A, clearing zones corresponding to the molecular weight of the enzyme were observed in the gels submerged in CaCl and MnCl and are indicated on the right. For ScrM, clearing zones were observed in all gels (although weaker in the presence of ZnCl), including the EDTA-treated control. Molecular weights (kDa) are indicated on the left, and the amount of protein loaded is indicated at the top of each image. The uncropped gel images are provided in Supplemental Fig. S8 (Zoocin A) and S9 (ScrM).

We assessed the long-term stability of Zoocin A and ScrM by storing enzyme stocks at room temperature (∼ 21 °C) for eight weeks and measuring their activity after 1 and 3 days and after 1, 2, 3, and 8 weeks. Both enzymes retained activity throughout the experiment, although activity decreased slightly after eight weeks (Fig. 6A). Subsequently, both enzymes were analysed by SDS-PAGE to determine whether the decrease in enzymatic activity was associated with protein degradation. Both samples showed bands corresponding to their molecular weights, with slightly reduced band intensity compared with the respective controls, suggesting that the decrease in activity was consistent with minor enzyme degradation (Fig. 6B).

**Fig. 6.**
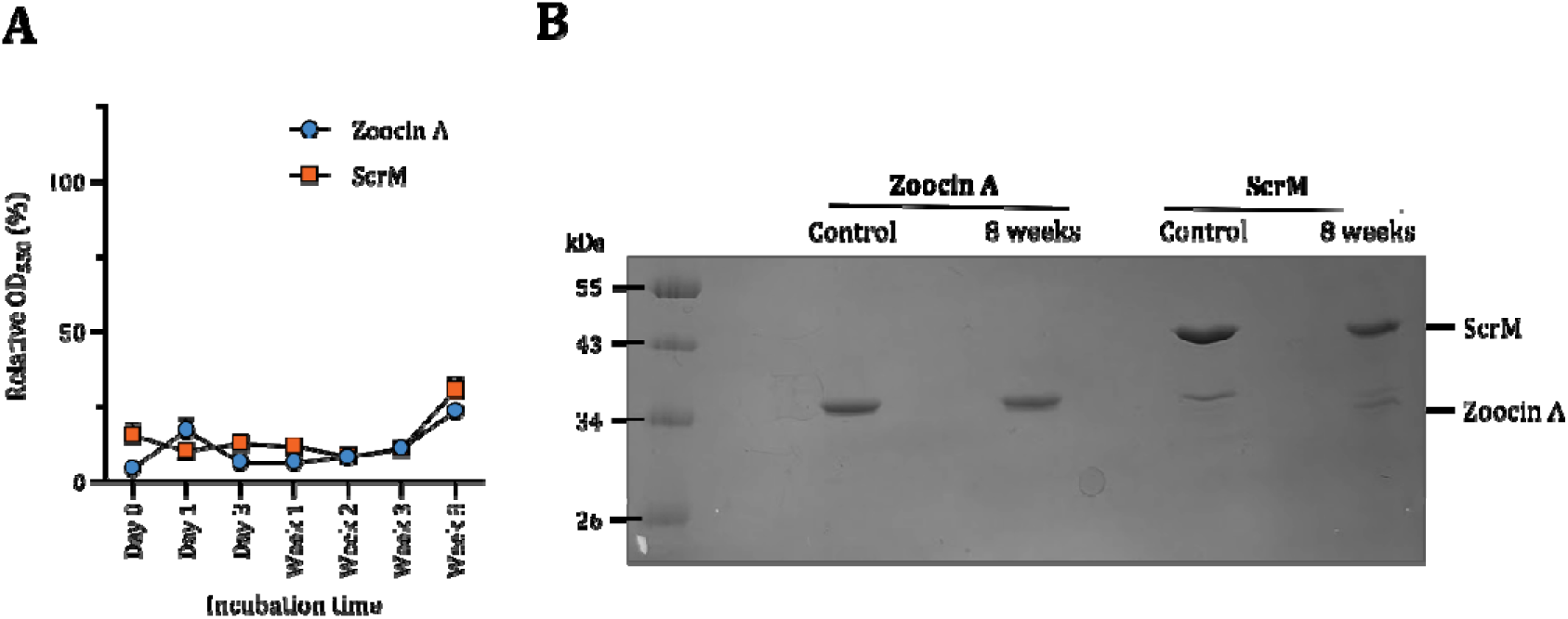
Long-term stability of Zoocin A and ScrM. Panel A: Zoocin A and ScrM were incubated at room temperature in dH_2_O for 8 weeks, and their activity was measured by adding them to cultures of *S. agalactiae* and *S. dysgalactiae*, respectively, at 0, 1, and 3 days, and 2, 3, and 8 weeks of incubation. Both enzymes were added to a final concentration of 2 µg/mL. The relative OD_550_ was calculated by comparing the OD_550_ of the treated cultures 1 hour after enzyme addition to that of the corresponding untreated control. Panel B: SDS-PAGE analysis of Zoocin A and ScrM after 8 weeks of incubation at room temperature to assess protein degradation. A total of 1.5 µg of each enzyme was loaded onto the gel. The bands corresponding to Zoocin A (31 kDa) and ScrM (41 kDa) are indicated on the right and showed slightly lower intensity after 8 weeks of incubation compared with the controls. Molecular weights of the protein standard are indicated on the left.

### Zoocin A and ScrM bind to and cause lysis at the division site

Previous work on ScrM demonstrated that this enzyme localises to the septal region of susceptible cells and that the binding was predominantly dependent on the conserved C-terminal domain (CCD) (Mårli et al., 2025). To investigate the binding pattern of Zoocin A, we purified the target recognition domain (TRD) of Zoocin A fused to sfGFP and added it to cultures of both Zoocin-susceptible (*S. agalactiae*) and -resistant (*S. suis*) species. Fluorescence microscopy showed that TRD-sfGFP localised predominantly to the septal regions of *S. agalactiae* cells, indicating that, similarly to ScrM, Zoocin A targets the division site and that the TRD is sufficient for septal localisation (Fig. 7A and B). No sfGFP signal was detected on the surface of the resistant *S. suis* cells (Fig. 7A).

**Fig. 7.**
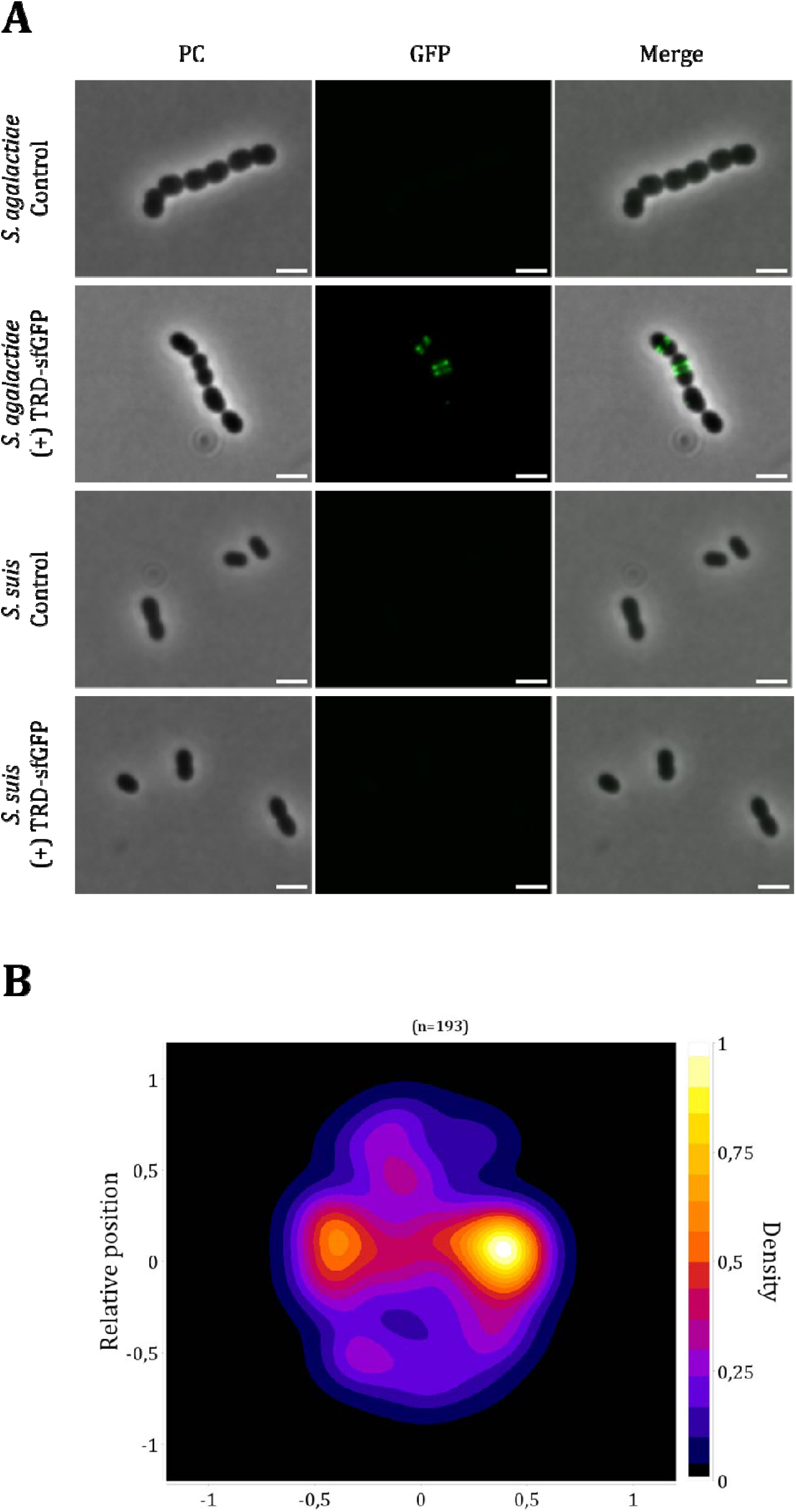
Binding of Zoocin A(TRD)-sfGFP to *S. agalactiae* and *S. suis*. Panel A: Phase-contrast (PC), GFP fluorescence (GFP), and merged (Merge) images of *S. agalactiae* and *S. suis* cells incubated with (+) or without (control) Zoocin A(TRD)-sfGFP. GFP fluorescence was observed in the septal regions of *S. agalactiae* cells, whereas no fluorescence was detected on *S. suis* cells. Images were processed using ImageJ (Rueden et al., 2017). Scale bars represent 2 µm. Panel B: Heatmap analysis showing the binding pattern of Zoocin A(TRD)-sfGFP on *S. agalactiae* cells. The analysis was performed using the MicrobeJ plugin in ImageJ (Ducret et al., 2016; Rueden et al., 2017). A total of 193 cells were analysed.

Following the observation that Zoocin A and ScrM localise to the septal region, we used transmission electron microscopy (TEM) to determine whether this localisation corresponds to the site of cell wall degradation. Among cells treated with Zoocin A and ScrM, we frequently observed lysed cells, characterised by the complete loss of intracellular contents (Fig. 8). The sites of cell rupture were identified as distinct openings in the septal regions of these cells. This indicates that enzymatic activity is directed toward the division site where the enzymes are enriched on the cell surface. Most affected cells appeared to be undergoing division, suggesting that septal cleavage may be associated with actively dividing cells.

**Fig. 8.**
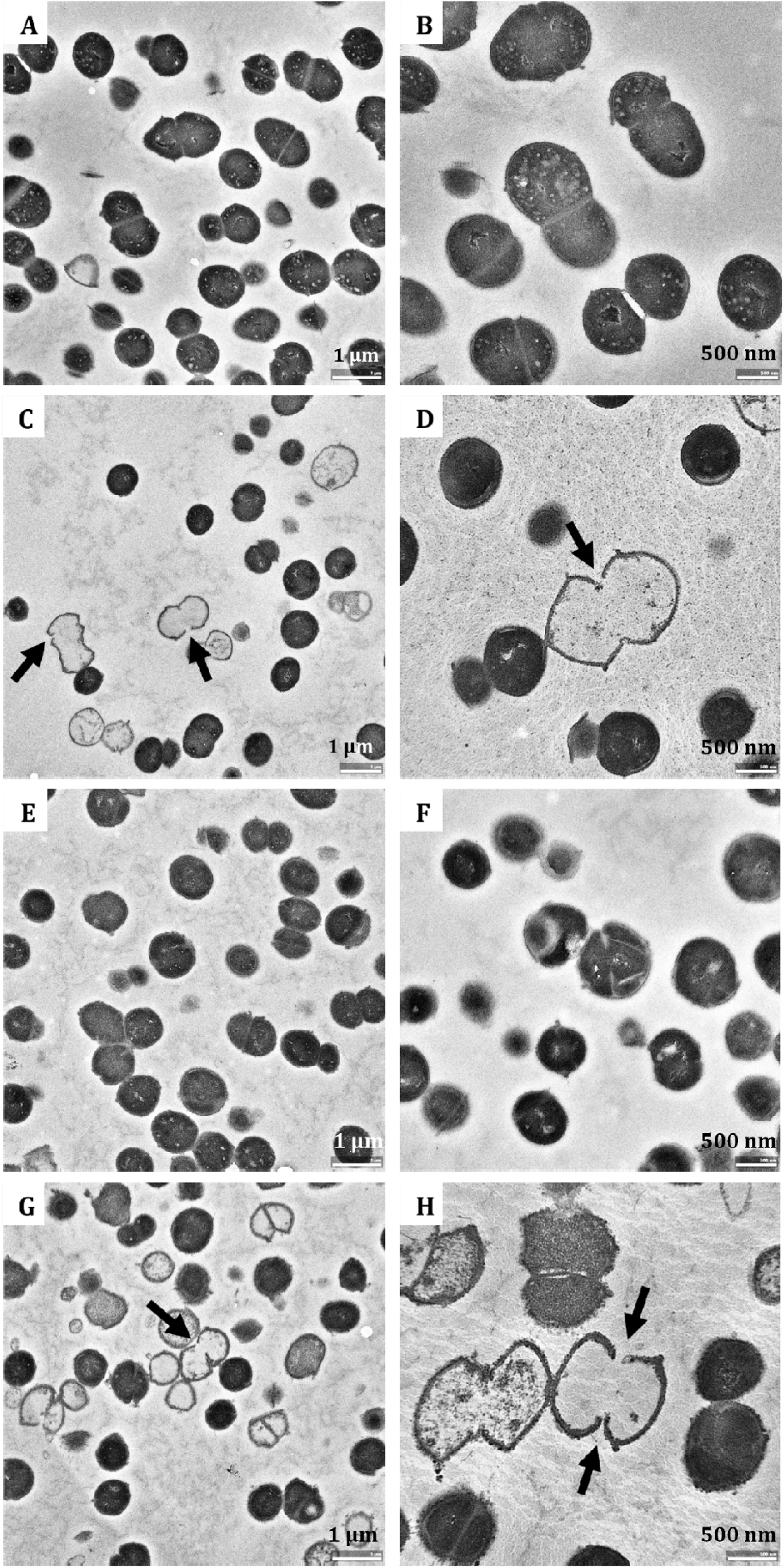
Transmission electron micrographs of Zoocin A-treated *S. agalactiae* and ScrM-treated *S. dysgalactiae*. Panels A and B: untreated controls of *S. agalactiae* cells. Panels C and D: *S. agalactiae* cells incubated with 4 µg/mL Zoocin A. Panels E and F: untreated controls of *S. dysgalactiae* cells. Panels G and H: *S. dysgalactiae* cells incubated with 4 µg/mL ScrM. Enzyme-treated cells (C, D, G, and H) showed loss of cellular contents compared with the respective untreated controls (A, B, E, and F). Openings in the septal regions of empty cells are indicated by arrows.

### Effects of Zoocin A and ScrM on *S. mutans* biofilms

A major medical challenge is that pathogens can form biofilms that protect bacteria from disinfectants and antibiotics. Because Zoocin A showed significant lytic activity against planktonic bacteria, we tested its ability to disrupt *S. mutans* biofilms, the main causative agent of dental caries. By using a biofilm-oriented antiseptics test (BOAT) (Junka et al., 2014), which measures the metabolic activity in biofilms, we observed a 63% average reduction in metabolically active cells in established biofilms at the highest concentration (200 µg/mL) of Zoocin A (Fig. 9A). We then included ScrM to determine whether combining the two enzymes could enhance the reduction. On its own, ScrM caused no detectable reduction (Fig. 9B), but when the biofilms were treated with both enzymes (200 µg/mL of each), metabolically active cells decreased by 83.8% on average, suggesting a synergistic or additive effect between the two enzymes (Fig. 9C). Next, we wanted to explore if the enzymes could act preventively against biofilm formation. The aim was to assess whether the enzymes could be used to coat a plastic surface in a dried form and subsequently prevent biofilm development upon inoculation with *S. mutans* cells. To determine whether drying affected enzyme activity, Zoocin A and ScrM were vacuum-dried, resuspended in water, and assessed for activity against planktonic cells. Following rehydration, Zoocin A retained full activity, whereas ScrM could not be dissolved in water and exhibited a total loss of activity (Supplemental Fig. S10). Based on this, Zoocin A was evaluated for its ability to prevent biofilm formation when applied as a dry coating. Enzyme suspensions were added to microtiter wells and dried at 37°C prior to inoculation with *S. mutans* cells. Surprisingly, biofilm formation was not reduced by the dried enzyme coating, despite the enzyme retaining activity against planktonic cells (Fig. 9D).

**Fig. 9.**
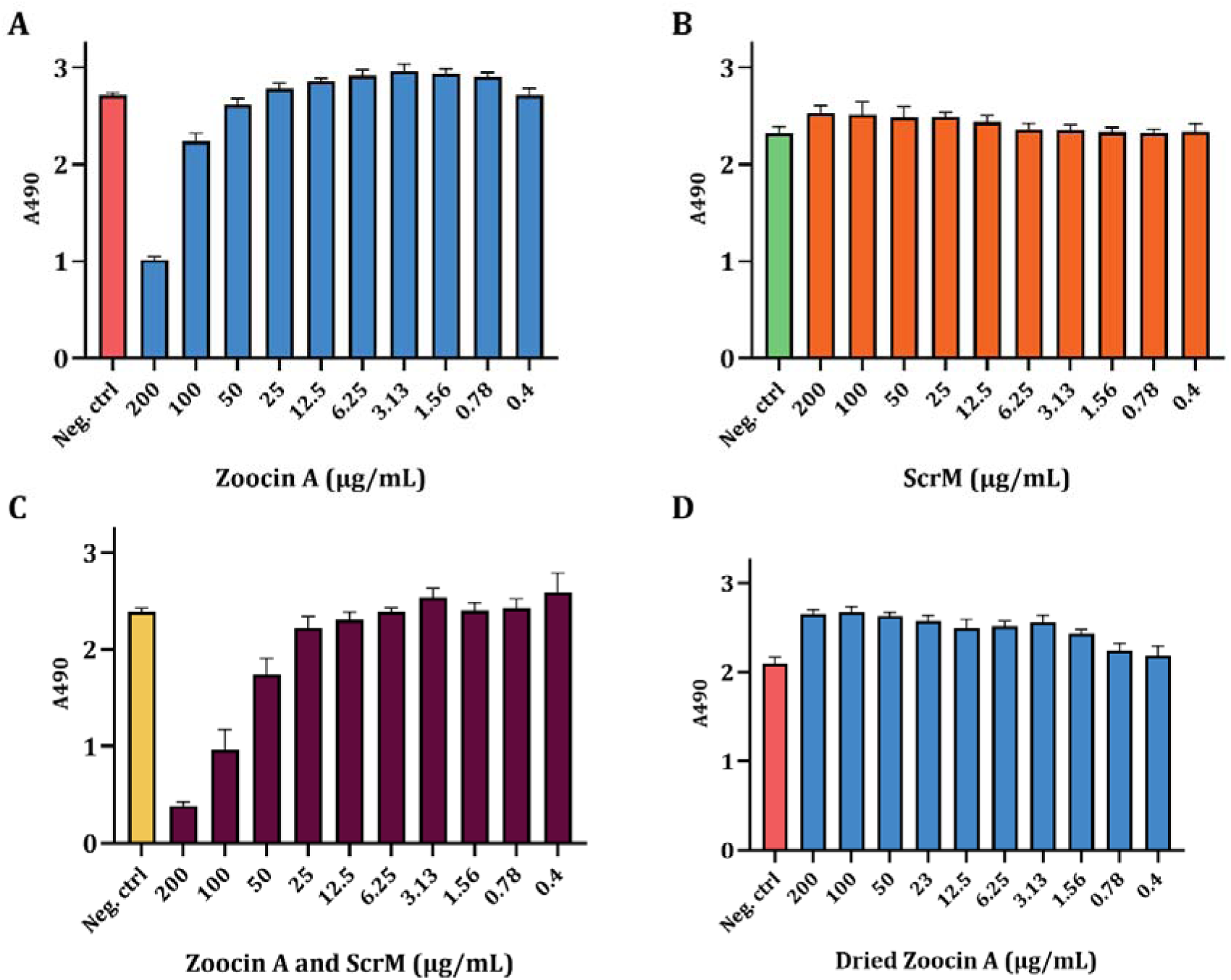
The activity of Zoocin A and ScrM against *S. mutans* biofilms using the biofilm-oriented antiseptics test (BOAT). Enzymes were added at concentrations ranging from 0.4 to 200 µg/mL to 24-hour-old biofilms of *S. mutans.* Triphenyl tetrazolium chloride (TCC), an indicator of cellular respiration, was added. Dehydrogenases in metabolically active cells reduce colourless TCC to red triphenylformazan (TPF), which can be measured at 490 nm. Untreated biofilms were included as negative controls. Panel A: *S. mutans* biofilms treated with Zoocin A. Panel B: *S. mutans* biofilms treated with ScrM. Panel C: *S. mutans* biofilms treated with both Zoocin A and ScrM. Panel D: *S. mutans* biofilms in which the inoculum culture was added to wells containing dried Zoocin A.

## Discussion

Previous studies of CbpD from *S. pneumoniae* and *S. mitis* have shown that it is a highly efficient lytic enzyme (Eldholm et al., 2010; Johnsborg et al., 2008; Straume et al., 2020), suggesting that fratricins can function as effective antimicrobial enzymes that combine potent bacteriolytic activity with a high degree of target specificity. We selected nine streptococcal species to cover all streptococcal phylogenetic groups and include all domain structure types of putative fratricins. Of these, we successfully purified four candidates using the chosen method (Supplemental Fig. S1). Zoocin A (*S. agalactiae*), ScrM (*S. dysgalactiae*), and CbpD-Sv (*S. vestibularis*) were investigated for antimicrobial activity and range, and Zoocin A and ScrM were selected for further activity and stability characterisation.

Among the putative fratricins investigated in this study, LytF proteins proved challenging to evaluate as antimicrobial candidates since none were successfully expressed and purified using *E. coli* as the host. Published data show that LytF proteins are indeed muralytic enzymes (Berg, Ohnstad, et al., 2012; Cullin et al., 2017; Moe et al., 2026), however, the capacity of LytF proteins to lyse target cells seems to vary between species. For example, in *S. mutans*, LytF has been reported to function as an autolytic enzyme possibly involved in fratricide (Eaton & Jacques, 2010). LytF from *S. sanguinis*, on the other hand, does not exhibit autolytic activity and therefore does not appear to function as a fratricin but instead contributes to natural transformation by facilitating extrusion of the competence pilus (Moe et al., 2026). Together, these observations suggest that LytF proteins may require individual functional validation before being considered as antimicrobial candidates. Additionally, because LytF has been shown to lyse *E. coli*, an alternative host for recombinant expression is needed to purify the protein in sufficient amounts (Dufour & Lévesque, 2013). An ideal candidate for an antimicrobial protein should be relatively simple to purify, and this is currently not the case for LytF proteins (Cullin et al., 2017; Moe et al., 2026).

Zoocin A, ScrM and CbpD-Sv displayed distinct antimicrobial spectra, although susceptibility patterns broadly followed streptococcal phylogenetic relationships. Zoocin A exhibited the broadest target range, with activity against members of all investigated phylogenetic groups except the suis group (Table 1). This broad activity profile differs from the classical concept of fratricins as more narrowly targeting enzymes. Among the tested streptococci, only members of the mitis and suis groups were resistant to Zoocin A, highlighting its potential as a targeted antimicrobial enzyme with activity against a wide range of streptococcal pathogens. Outside the *Streptococcus* genus, only *E. faecalis* displayed (partial) susceptibility to Zoocin A, while *Enterococcus faecium* was resistant (Table 1). As enterococci are closely related to streptococci and were historically classified within the same taxonomic group, the limited activity against *E. faecalis* is consistent with some conservation of target features between these genera. Thus, Zoocin A appears to retain a predominantly *Streptococcus*-targeted activity profile, suggesting that its use as an antimicrobial enzyme may result in limited activity against non-target bacterial taxa. In agreement with these findings, a Zoocin A homologue, SALys78925, was reported to be active against pyogenic group streptococci and *E. faecalis*, but not *Staphylococcus* (Yoon et al., 2025). Compared with Zoocin A, ScrM and CbpD-Sv displayed narrower antimicrobial spectra. This specificity may be advantageous for developing targeted antimicrobial strategies. Both enzymes were active against all three tested members of the pyogenic group, including *S. agalactiae*, *S. dysgalactiae*, and *S. pyogenes* (Table 1). These species are clinically relevant pathogens in both human and veterinary medicine. While beta-lactam resistance remains rare among these organisms, recent reports of reduced susceptibility in *S. pyogenes* highlight the importance of continued efforts to identify alternative antimicrobial strategies (Amusin Daniel et al., 2026; Musser James et al., 2020; Vannice et al., 2020). Furthermore, *S. agalactiae* and *S. dysgalactiae* are important pathogens in aquaculture and livestock production, respectively. Targeted enzymes such as ScrM and CbpD-Sv may represent promising candidates for further investigation in settings where minimising conventional antibiotic use is a priority.

Zoocin A was found to bind to the septum of susceptible cells, and the binding was attributed to the TRD domain (Fig. 7). The CCD domain has likewise been shown to be responsible for the same binding pattern for ScrM (Mårli et al., 2025). Both Zoocin A and ScrM were also observed to cause lysis by attacking the cell wall at the septal region, in line with the notion that fratricins specifically target the cell wall at the division zone (Fig. 8). Many of the lysed cells appeared to be undergoing division, although this was not quantified. These observations suggest that the enzymes preferentially target cells undergoing division. Alternatively, the enzymes may cleave the cell wall more broadly, but the specific enrichment of enzymes binding to the division zone results in more cell wall damage at this site, eventually leading to cell rupture. Although not yet fully understood, the septum is known to be sensitive to peptidoglycan hydrolases and CbpD-Sp, and several putative fratricins exhibit specific binding to the division zone of target cells (Berg, Ohnstad, et al., 2012; Eldholm et al., 2010; Mårli et al., 2025). The TRD of Zoocin A was unable to bind to *S. suis*, which is completely resistant to Zoocin A (Fig. 7). This is consistent with findings from the Zoocin A homologue encoded by *Streptococcus equi* subsp. *zooepidermicus* strain 4881 (hereafter Zoocin A-4881), where susceptibility is largely correlated with binding (Akesson et al., 2007). The metal-cation dependence observed for Zoocin A in this study is consistent with that reported for the closely related homologue SALys78925 (Yoon et al., 2025). In our experiments, we detected muralytic activity in the presence of Mn² and Ca², but not Zn². Likewise, addition of Mn^2+^ was shown to increase the activity of SALys78925, while Zn^2+^ reduced it by approximately 80% (Yoon et al., 2025). M23 metallopeptidases are zinc-dependent and Zoocin A-4881 has been reported to depend primarily on Zn^2+^ (Chen et al., 2008; Grabowska et al., 2015; Razew et al., 2022). The absence of activity following Zn^2+^-addition to Zoocin A may reflect inhibition by excess Zn^2+^ rather than a lack of zinc dependence, as additional Zn^2+^ binding has been shown to inhibit the activity of lysostaphin, another M23 metallopeptidase (Ke et al., 2021).

Zoocin A and ScrM displayed a similar and expected level of stability at different pH values and at high temperatures. Activity was largely retained under neutral to slightly alkaline conditions but declined outside this range (Fig. 4), and both enzymes were inactivated at relatively moderate temperatures (Fig. 2). More notably, both enzymes remained active at relatively high NaCl concentrations (Fig. 3) and after storage at room temperature for 8 weeks (Fig. 6). In particular, ScrM retained nearly full activity with up to 0.5 M added NaCl and remained partially active even at 1 M (Fig. 3). The effect of NaCl on enzyme activity may be better measured using cells naturally suited to high-salt environments, as reduced osmotic pressure in the *Streptococcus* cells may prevent lysis despite continued antimicrobial activity. At NaCl concentrations of 0.5 M and above, the growth curves (Supplemental Fig. S5 and S6) show that although lytic activity is reduced compared with lower salt concentrations, growth is completely halted, suggesting the cells may be dead. We therefore speculate that both Zoocin A and ScrM may retain substantial activity at NaCl concentrations above 0.5 M, but that reduced osmotic pressure in the cells is preventing lysis. The long-term stability of the enzymes at room temperature may be advantageous in applications by simplifying storage and handling. However, this experiment was conducted with enzymes in dH_2_O, and the observed stability may vary under alternative storage conditions or in different formulations. Although the stability and activity observed under these experimental conditions may differ from those *in vivo*, these results indicate that Zoocin A and ScrM retain activity under conditions broadly representative of potential applications.

Zoocin A reduced *S. mutans* biofilm formation at the highest concentrations tested, whereas ScrM had no detectable effect on these biofilms (Fig. 9A and B). As Zoocin A is considerably smaller than ScrM, its size may better facilitate diffusion through the extracellular polymeric matrix of biofilms, thereby improving access to bacterial cells. A slight synergistic or additive effect was observed when the two enzymes were combined (Fig. 9C). Previous studies have shown that Zoocin A-4881 is effective against *S. mutans* biofilms when combined with lauricidin (Lester & Simmonds, 2012). Similarly, Zoocin A (*S. agalactiae*) and possibly ScrM may exhibit enhanced activity when used in combination with biofilm-disrupting agents. Although Zoocin A retained its activity after drying and resuspension (Supplemental Fig. S10A), we were surprised to find that dried Zoocin A did not reduce the formation of new biofilms on surfaces coated with the enzyme (Fig. 9D). This approach was based on the potential use of Zoocin A as a preventive surface coating to inhibit microbial colonisation and thereby reduce the risk of infection, but it likely requires further optimisation. Nevertheless, Zoocin A’s ability to retain activity following drying and resuspension may be advantageous for other applications and could simplify storage and handling.

In this study, we investigated putative fratricins as potential antimicrobial enzymes and successfully purified Zoocin A, ScrM, and CbpD-Sv. All three enzymes were predominantly active against *Streptococcus* species, with the specific species targeted varying somewhat between them. Zoocin A and ScrM retained activity under a range of relevant conditions, showed particularly high tolerance to NaCl, and remained stable for several weeks to months at room temperature. Furthermore, we identified the site of cell wall cleavage to be concentrated at the septal region of susceptible cells. Although Zoocin A exhibited activity against *S. mutans*, its application against biofilms needs further optimisation. Collectively, our findings suggest that fratricins are attractive candidates for developing novel antimicrobial formulations to prevent certain streptococci from colonising the host.

## Methods

### Bacterial species, strains, and growth conditions

Bacterial species and strains used in this study are listed in Supplemental Table S1. All liquid streptococcal cultures were grown statically in C-medium at 37 °C, unless otherwise specified (Lacks & Hotchkiss, 1960). For plate cultivation, bacteria were grown on Todd Hewitt (TH) agar under anaerobic conditions (<1% O_2_) obtained using AnaeoroGen Satchets from Oxoid in sealed containers. The *E. coli* strains were grown aerobically in lysogeny broth (LB) or lysogeny agar (LA) at 37 °C. For plasmid retention in *E. coli*, the growth medium was supplemented with 100 µg/mL ampicillin.

### Plasmid construction and transformation

The primers used in this work are listed in Table S2. The plasmids for overexpression of putative fratricins were constructed by amplifying the native putative fratricin gene (without its signal sequence) using a forward primer with a 5’ end containing a start codon (CATATG, restriction site for NdeI), followed by a sequence encoding a 6x His-tag and a TEV protease proteolytic cleavage site and a reverse primer containing the restriction site for HindIII. The resulting PCR product was cut with restriction enzymes NdeI and HindIII and ligated into the plasmid pRSET A. Finally, the ligation reaction was transformed into chemically competent *E. coli* cells by heat shock at 42 °C. Transformants were screened by colony PCR, and the plasmid sequence was confirmed by Sanger sequencing (GATC, Eurofins).

Plasmid pRM42 (pRSET A-*sfGFP*-*zoocin*(TRD)) was constructed using the same approach, but *sfGFP* and *zoocin*(TRD) were fused using overlap extension PCR (OE-PCR) prior to NdeI and HindIII cleavage of the resulting fragment.

### Purification of putative fratricins

The putative fratricins were overexpressed in 1000 mL cultures of *E. coli* BL21 (RM62, RM64-67) carrying plasmids encoding the respective fratricin genes. Expression was induced at an OD_600_ of 0.4 using 0.1 mM IPTG and carried out at 25 °C for 4 hours. The cells were harvested at 12,000 g for 5 minutes, and the pellet was resuspended in 20 mL 10 mM Tris-HCl, 500 mM NaCl, 20 mM imidazole before being lysed using a French Pressure cell system (Homogenising Systems Ltd.). Insoluble cell components were removed by centrifugation at 15,000 g for 5 minutes at 15 °C, and the protein-containing supernatant was filtered through a 0.45 µm filter. The filtrate was then passed through a 1 mL HisTrap HP column (Cytiva Life Sciences) using the Äkta pure system (Cytiva). The column was washed with 10 column volumes of 10 mM Tris-HCl (pH 7.4), 500 mM NaCl, and 20 mM imidazole, and the putative fratricins were eluted using a linear gradient of imidazole from 20 to 500 mM under the same buffer conditions. Finally, the eluate was dialysed using a buffer of 10 mM Tris-HCl, 150 mM NaCl, pH 7.4, for 1 hour at room temperature. The presence and purity of the proteins were assessed by SDS-PAGE (Supplemental Fig. S1) before storage at −80°C.

The fluorescently labelled target recognition domain of Zoocin A (sfGFP-Zoocin(TRD)) was overexpressed from a 50 mL culture of strain RM135 following the same principle described above and purified using a Protino Ni-TED (Macherey-Nagel) kit.

### Determining MIC_50_

The concentrations of Zoocin A and ScrM required to lyse approximately half of *S. agalactiae* and *S. dysgalactiae* cultures, respectively, were determined by titration. Two-fold dilution series of enzymes were added to actively growing cultures at OD_550_ = 0.2 resulting in final concentration of enzymes ranging from 30 – 0.06 µg/mL. The experiment was performed using 96-well microtiter plates that were incubated at 37 °C in a SYNERGY H1 microplate reader. OD_550_ was monitored automatically every 5 minutes, with orbital shaking at 300 rpm prior to each measurement.

### Stability assays

For all assays described below *S. agalactiae* (RM22) and *S. dysgalactiae* (MA201) were grown to OD_550_ = 0.2-0.4, diluted to OD_550_ = 0.05, and grown further to reach OD_550_ = 0.2. Unless otherwise stated, OD_550_ was measured every 5 min using a SYNERGY H1 microplate reader. In all assays, *S. agalactiae* was used as an indicator for Zoocin A and *S. dysgalactiae* for ScrM. The enzymes were added when the cultures reached an OD_550_ of approximately 0.2, at a concentration corresponding to 50% survival of the control cultures. Because there were minor variations in the survival response, likely due to differences in experimental setup between assays, enzyme concentrations were adjusted when necessary. Assays were initiated at the predetermined MIC_50_ (approximately 1 µg/mL for Zoocin A and 2 µg/mL for ScrM) and increased to 2–4 µg/mL when required to obtain a survival response closer to 50%. In all experiments, non-treated parallels of the cultures were included as a control.

#### Heat stability

The heat stability of Zoocin A and ScrM was determined by incubating the enzymes for 1 hr at 37 °C and 45 °C, then briefly cooling them to 4 °C before adding them to the cultures at a final concentration of 2 µg/mL. Controls included an untreated culture and a culture treated with enzymes incubated at 4 °C.

#### Activity in PBS and TBS

Cells in cultures with an OD_550_ = 0.4 were harvested by centrifugation at 16,000 *g* for 1 minute, washed with either PBS or TBS and resuspended in the corresponding buffer to the original culture volume. The cell suspensions were transferred to a 96-well plate, and the OD_550_ was measured every five minutes. Zoocin A and ScrM were added after approximately 15 minutes to a final concentration of 2 µg/mL. The relative OD_550_ was determined by comparing the OD_550_ measured before enzyme treatment to that measured 1 hour after.

#### NaCl tolerance

When cultures reached an OD_550_ of approximately 0.2, NaCl was added at concentrations of 0.05, 0.15, 0.25, 0.5, and 1 M. C-medium was used for these assays, as the enzymes showed reduced activity in PBS or TBS (Supplemental Fig. S3). Enzyme activity was assessed after addition of enzymes pre-incubated at the respective NaCl concentrations to a final concentration of 4 µg/mL. As controls, cultures without additional NaCl were treated with enzymes pre-incubated in dH_2_O.

#### pH range assay

Cultures of *S. agalactiae* and *S. dysgalactiae* were grown to an OD_550_ of 0.2-0.4 before the cells were harvested and resuspended in pH-adjusted C-medium (pH 3-10). The OD_550_ of the cultures was measured before and 5 minutes after adding Zoocin A and ScrM to a final concentration of 4 µg/mL. Cultures suspended in pH-adjusted medium without enzymes were included as negative controls.

#### Vacuum-drying

Solutions of Zoocin A and ScrM at 0.5 mg/mL were vacuum-dried using a Savant^TM^ Speed Vac Concentrator (Thermo Fisher) for 1 hour, then resuspended in dH_2_O before being added to the growing cultures at MIC_50_ concentrations.

#### Chelating agents and divalent cations

The effect of chelating agents on Zoocin A and ScrM was assessed by adding ethylenediaminetetraacetic acid (EDTA) to both enzyme solutions and bacterial culture to a final concentration of 1 mM, 30 minutes before the enzymes were added at MIC_50_ concentrations. Zymogram analyses were conducted based on the protocol described by Leclerc and Asselin (1989), with minor modifications. The substrate cultures (*S. agalactiae* RM22 for Zoocin A and *S. dysgalactiae* MA201 for ScrM) were grown to OD_550_ = 0.8 in 100 mL. Cells were collected by centrifugation at 5,000 × *g*, washed once with deionised water, and resuspended in 1.25 mL of 1.5 M Tris-HCl (pH 8.8) before heat treatment at 100 °C for 10 minutes. SDS-PAGE gels were cast by incorporating the substrate suspension in the 12% resolving gel. A 4% stacking gel was cast on top. Purified Zoocin A and ScrM were diluted twofold, mixed with a 2X SDS sample buffer (0.125 M Tris-HCl, 4% SDS, 0.30 M β-2-mercaptoethanol, 20% glycerol, and 0.01% bromophenol blue) and heat-treated at 95 °C for minutes. Following electrophoresis, the gels were submerged in a refolding buffer (100 mM NaCl, 0.5% Triton X 100, 20 mM Tris HCl, pH 7.4) containing 1 mM EDTA (ethylenediaminetetraacetic acid) for 30 minutes. Subsequently, the gels were washed in deionised water for 2 x 15 minutes before being submerged in refolding buffers containing 1 mM of either MnCl_2_, ZnCl_2_, or CaCl_2_ until clearing zones appeared. A gel submerged in deionised water served as a negative control.

#### Long-term stability at room temperature

To determine the time frame during which Zoocin A and ScrM remain stable and active at ambient temperature, the enzymes were stored at room temperature (∼21 °C) and added to the cultures after 1 and 3 days, and after 1, 2, 3, and 8 weeks to a final concentration of 2 µg/mL. After 8 weeks, protein degradation was assessed by SDS-PAGE. A total of 1.5 µg of each enzyme was loaded onto the gel, and the intensity of the bands was compared with that of corresponding controls kept at −20 °C.

### Target range assays

Overnight cultures of the target species were diluted to an OD_550_ of 0.05, incubated at 37 °C until they reached an OD_550_ of 0.2-0.4, and diluted again to an OD_550_ of 0.025-0.05 in C-medium. Streptococcal species were grown in C medium, and non-streptococcal species were grown in BHI. The OD_550_ of the cultures was measured every 5 minutes using a SYNERGY H1 plate reader, and Zoocin A, ScrM, and CbpD-Sv were added at OD_550_ ∼ 0.2 to a concentration of 30 µg/mL.

### Transmission Electron Microscopy (TEM)

Cultures *of S. agalactiae* (RM22) and *S. dysgalactiae* (MA201) (10 mL each) were grown to an OD_550_ of 0.2 before the cells were harvested by centrifugation at 5,000 *g* for 5 minutes. The cell pellets were resuspended in 1 mL C-medium, and Zoocin A and ScrM were added (4 µg/mL). Non-treated cultures of both species were included as negative controls. One minute after enzyme addition, the cells were fixed using a 2.5% glutaraldehyde, 4% paraformaldehyde, 0.05 M cacodylate buffer (pH 7.4) for 1 hour at room temperature, followed by an overnight incubation at 4°C. The cells were then harvested by centrifugation at 8,000 g and washed three times with a 0.1 M cacodylate buffer before being embedded in a 3% low-melt agarose (Bio-Rad) gel. Excess agarose was removed, and the cells were contrast-stained with 1% osmium in 0.1 M cacodylate buffer for 1 hour in darkness. The cells were then washed with 0.1 M cacodylate buffer and dehydrated through graded acetone increments (50-100%): three washes in 50%, two 10-minute washes in 70% and 90%, and three 20-minute washes in 100% acetone. Finally, the pellets were embedded in EPON Epoxy Resin, shaved to ∼ 60 nm-thick sheets using a Leica EM UC6 Ultramicrotome, and imaged using a JEM 2100-Plus scanning transmission electron microscope (Jeol).

### Fluorescence microscopy

Cultures of *S. agalactiae* (RM22) and *S. suis* (RM35) were grown to an OD_550_ of 0.2, and sfGFP-Zoocin(TRD) was added (8 µg/mL). Following a 10-minute incubation at 37°C, the cells were harvested by centrifugation at 16,000 *g* for 1 minute and resuspended in half the original culture volume of PBS. The cells were then deposited onto a microscopy slide covered with a thin layer of 1.2% agarose. Images were taken with a Zeiss LSM700 microscope, and the ImageJ plugin MicrobeJ was used for image analysis (Ducret et al., 2016; Rueden et al., 2017).

### Biofilm-oriented antiseptics test (BOAT)

The BOAT assay was performed as described by Junka et al. (2013) with minor modifications. Overnight cultures of *S. mutans* were diluted 1:100 in BHI supplemented with 1% sucrose, and 100 µL was added to each well of a 96-well microtiter plate. Wells containing only BHI with sucrose served as negative controls. The plate was incubated anaerobically at 37°C for 24 h to allow biofilm formation. Following incubation, the biofilms were washed with 0.9% NaCl to remove planktonic cells. Subsequently, 150 µL of a two-fold serial dilution of the enzymes (final concentrations ranging from 200 to 0.4 µg/mL) was added to the wells, with each concentration applied to all wells within a single column. A column receiving no enzyme treatment served as a control. The plate was then incubated anaerobically at 37°C for 2 h. After treatment, the supernatant was removed, the biofilms were washed twice with 0.9% NaCl before 100 µL triphenyl tetrazolium chloride (TTC; 0.025% in BHI) was added, and the plate was incubated at 37°C for 5 h. TTC serves as an indicator of cellular respiration, as dehydrogenases in metabolically active cells reduce the colourless TTC to red triphenylformazan (TPF), which can be quantified by measuring absorbance at 490 nm. The supernatant was removed, and 200 µL of a 7:3 ethanol–acetone solution was added to each well to solubilise the triphenylformazan formed within the biofilms. Finally, the plate was incubated overnight at 37°C, and absorbance was measured at 490 nm the following day.

The enzyme-coated plastics assay was performed as described above, except that the wells were pre-coated with Zoocin A prior to inoculation by adding a solution of the enzyme in dH_2_O and incubating at 37°C until completely dry.

## Supporting information

Supplemental Sequences

## Acknowledgements

The work was funded by the Norwegian University of Life Sciences. We thank Lene Cecilie Hermansen at the Imaging Centre at the Norwegian University of Life Sciences for technical assistance with transmission electron microscopy analyses.

