## Supplemental figures and tables for "Exploring the antimicrobial potential of streptococcal fratricins"

**Supplemental tables**

**Table S1** Bacterial species and strains used in this study.

| **Species** | **Strain** | **Relevant characteristics** | **Source or reference** |
| --- | --- | --- | --- |
| *Streptococcus* strains | | | |
| *S. agalactiae* | RM22, NCTC 8181 |  | National Collection of Type Cultures |
| *S. anginosus* | NCTC 10713 |  |  |
| *S. cristatus* | NCTC 12479 |  |  |
| *S. dysgalactiae* | MA201 |  | (Porcellato et al., 2021) |
| *S. equinus* | ATCC 9812 |  | American Type Culture Collection |
| *S. mitis* | B6 |  | (Hakenbeck et al., 2012) |
| *S. mutans* | UA159 |  | American Type Culture Collection |
| *S. pneumoniae* | RH425 |  | (Johnsborg & Håvarstein, 2009) |
| *S. pyogenes* | iGAS83 |  | (Mårli et al., 2025) |
| *S. sanguinis* | SK36 |  | (Kilian et al., 2008) |
| *S. sobrinus* | ATCC 27352 |  | American Type Culture Collection |
| *S. suis* | 2019-40-11936 |  | Norwegian Veterinary Institute |
| *S. thermophilus* | BAA-250 |  | American Type Culture Collection |
| *S. vestibularis* | NCTC 12167 |  | National Collection of Type Cultures |
| *E. coli* strains | | | |
| *E. coli* | GeneHogs | Cloning host | Invitrogen |
| *E. coli* | BL21 | Expression host, F-, *ompT*, *hsdS*_B_ (r_B_- m_B_-), *gal*, *dcm* (DE3), pLys, CmR | Invitrogen |
| *E. coli* | RM44 | GeneHogs, pRSET A _6x_*his*-*TEV*-*zoocin A* (*S. agalactiae*) | This work |
| *E. coli* | RM45 | GeneHogs, pRSET A _6x_*his*-*TEV*-*lytF* (*S. anginosus*) |  |
| *E. coli* | RM46 | GeneHogs, pRSET A _6x_*his*-*TEV*-*lytF* (*S. equinus*) |  |
| *E. coli* | RM47 | GeneHogs, pRSET A _6x_*his*-*TEV*-*lytF* (*S. mutans*) |  |
| *E. coli* | RM48 | GeneHogs, pRSET A _6x_*his*-*TEV*-*fratricin* (*S. sobrinus*) |  |
| *E. coli* | RM49 | GeneHogs, pRSET A _6x_*his*-*TEV*-*lytF* (*S. vestibularis*) |  |
| *E. coli* | RM50 | GeneHogs, pRSET A _6x_*his*-*TEV*-*lytF* (*S. suis*) |  |
| *E. coli* | RM61 | BL21, pRSET A _6x_*his*-*TEV*-*zoocin A* (*S. agalactiae*) |  |
| *E. coli* | RM62 | BL21, pRSET A _6x_*his*-*TEV*-*lytF* (*S. anginosus*) |  |
| *E. coli* | RM64 | BL21, pRSET A _6x_*his*-*TEV*-*lytF* (*S. mutans*) |  |
| *E. coli* | RM65 | BL21, pRSET A _6x_*his*-*TEV*-*fratricin* (*S. sobrinus*) |  |
| *E. coli* | RM66 | BL21, pRSET A _6x_*his*-*TEV*-*lytF* (*S. vestibularis*) |  |
| *E. coli* | RM67 | BL21, pRSET A _6x_*his*-*TEV*-*crfP* (*S. suis*) |  |
| *E. coli* | RM135 | GeneHogs, pRSET A _6x_*his*-*sfGFP*-*zoocin A*(TRD) |  |
| *E. coli* | RM136 | BL21, pRSET A _6x_*his*-*sfGFP*-*zoocin A*(TRD) |  |
| Other species | | | |
| *Bacillus subtilis* | ATCC 6051 |  | American Type Culture Collection |
| *Enterococcus faecalis* | LMGT 3088 |  | LMGT* |
| *Enterococcus faecium* | LMGT 2763 |  |  |
| *Lactococcus garviae* | LMGT 3390 |  |  |
| *Lactococcus innocua* | LMGT 2785 |  |  |
| *Staphylococcus aureus* | NCTC 8325-4 |  | National Collection of Type Cultures |
| *Staphylococcus haemolyticus* | LMGT 4133 |  | LMGT* |

^*^ Laboratory of Microbial Gene Technology (LMGT), Norwegian University of Life Sciences, Ås, Norway

**Table S2** Primers used in this study.

| **Primer** | **Sequence (5’-3’)** | **Application** | **Source** |
| --- | --- | --- | --- |
| rm043 | GATCCATATGCATCATCATCATCATCATGAGAACCTGTACTTCCAAGGTGAAACAAATAATACTCAAAAAAGCTCAAA | pRSET A _6x_*his*-*TEV*-*lytF* (*S. anginosus*) | This work/Lab collection |
| rm044 | GATCAAGCTTTTATTTTGGATAAATGTAACTAACTGCTCC |  |  |
| rm045 | GCTACATATGCATCATCATCATCATCATGAGAACCTGTACTTCCAAGGTGATCAACAAAATGATGCACTGG | pRSET A _6x_*his*-*TEV*-*lytF* (*S. equinus*) |  |
| rm046 | GCATAAGCTTTTAATCTGGATAAATGTATGAAACATTTCC |  |  |
| rm047 | GCATCATATGCATCATCATCATCATCATGAGAACCTGTACTTCCAAGGTGATACAGAGGATAATCAGGTAACGATTAGC | pRSET A _6x_*his*-*TEV*-*lytF* (*S. mutans*) |  |
| rm048 | GCATAAGCTTTTAGTCAGGATAAATATAAGTTAATCTTCCCA |  |  |
| rm049 | ATCGCATATGCATCATCATCATCATCATGAGAACCTGTACTTCCAAGGTGATGAGCAGACGGCTACTAGTGG | pRSET A _6x_*his*-*TEV*-*fratricin* (*S. sobrinus*) |  |
| rm050 | GATCGAATTCTCAGTCCTTCTTAACTGGAACG |  |  |
| rm051 | CATATGCATCATCATCATCATCATGAGAACCTGTACTTCCAAGGTGATGAGTTGACCAACACCAATG | pRSET A _6x_*his*-*TEV*-*lytF* (*S. vestibularis*) |  |
| rm052 | GATCAAGCTTCTATGCTACTGCTACGCTGATATATCG |  |  |
| rm053 | GATCTCTAGATTAAGAAGGAGATATACATATGCATCATCATCATCATCATGAGAACCTGTACTTCCAAGGTGATACTTATGTCCGTCCAATTGA | pRSET A _6x_*his*-*TEV*-*zoocin A* (*S. agalactiae*) |  |
| rm054 | GATCAAGCTTCTATTTGTAAAGCAGTTCTTGACTGTTG |  |  |
| rm055 | GCTACATATGCATCATCATCATCATCATGAGAACCTGTACTTCCAAGGTAATGCACGAGGAGATGACTATCC | pRSET A _6x_*his*-*TEV*-*crfP* (*S. suis*) |  |
| rm056 | GCATAAGCTTTTAGGAGATGGCGATGTAGCG |  |  |
| rm084 | TGGCCGATTCATTAATGCAG | Sequencing of fratricin purification plasmids |  |
| rm085 | GAGTTGAGCTAAAAAAACACAACTATGA |  |  |
| rm086 | CGCTATTACGCCAGATCCG |  |  |
| rm087 | ACAGTGGGGAGCTGTTGC |  |  |
| ks11 | GATCCATATGCATCATCATCATCATCATAAACATCTTACCGGTTCTAAAGG | pRSET A *sfGFP*-*zoocin A*(TRD) |  |
| rm252 | TGCGGCCGCTCCACTAGTTT |  |  |
| rm279 | CTAGTGGAGCGGCCGCACCAGCTAACCCTAATTTTCAAAATG |  |  |

**Supplemental figures**

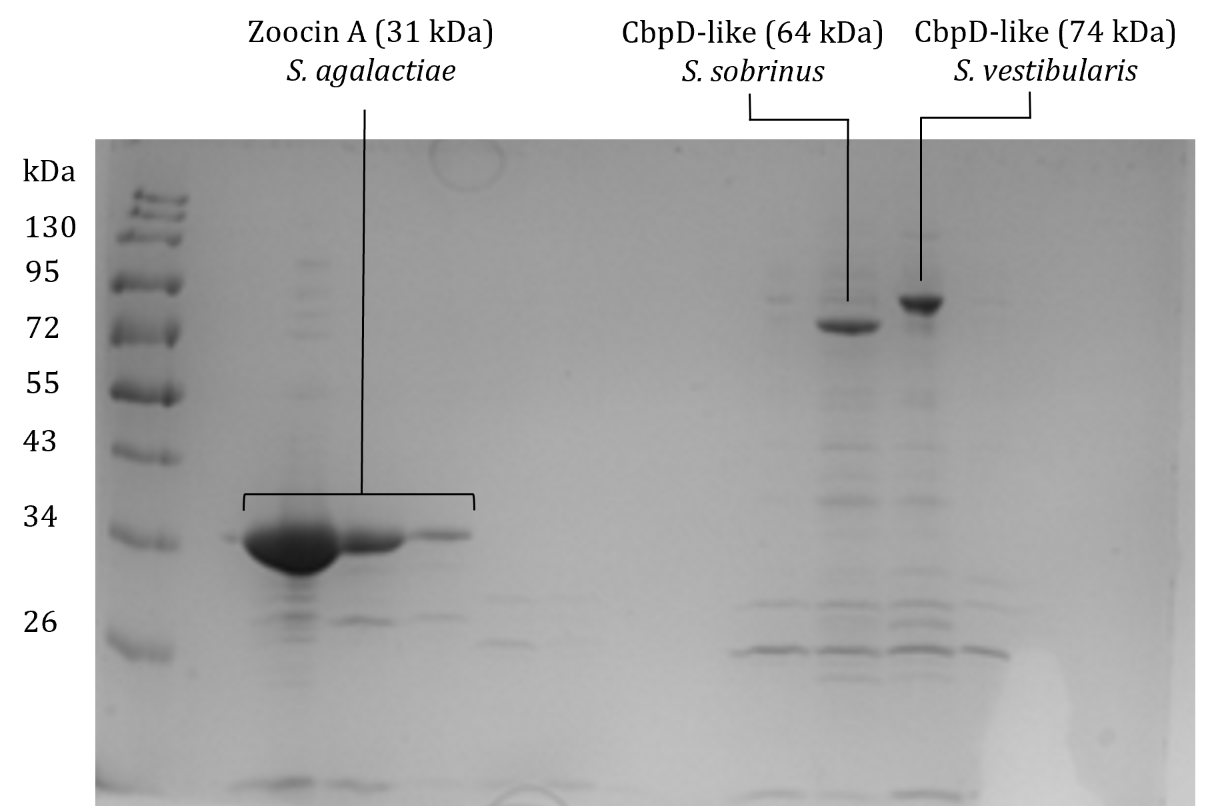

**Fig. S1** SDS-PAGE of immobilised metal affinity chromatography (IMAC) purified putative fratricins. The secretion signal sequences were replaced with histidine-encoding sequences, and the proteins were purified from *E. coli* BL21 transformed with the pRSET A vector carrying the genes encoding the proteins of interest. Protein bands corresponding to Zoocin A (31 kDa) from *S. agalactiae*, CbpD (64 kDa) from *S. sobrinus*, and CbpD (74 kDa) from *S. vestibularis* were visualised by Coomassie brilliant blue staining. Molecular weights (kDa) of the standard ladder are indicated on the left side of the image.

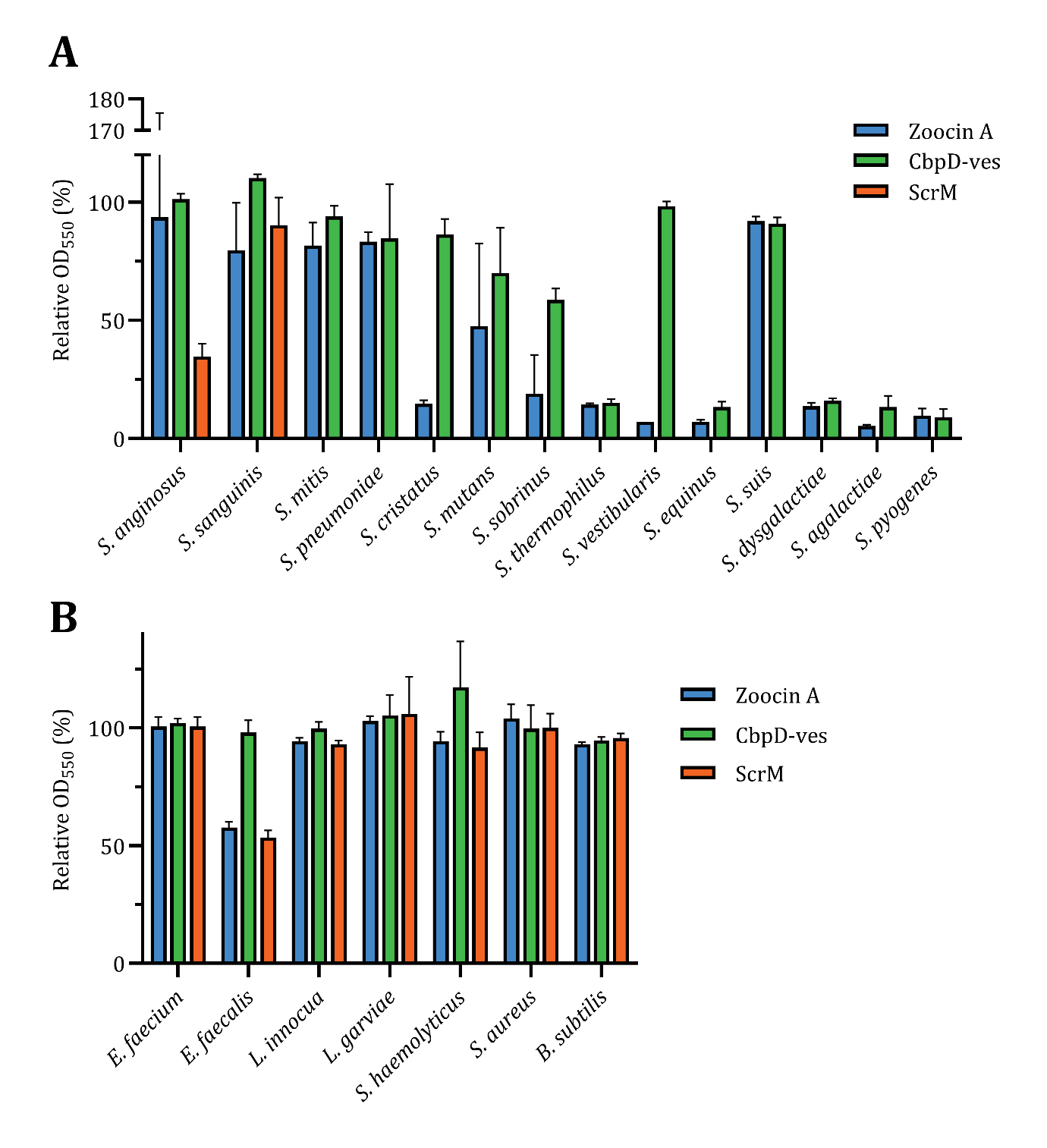

**Fig. S2** Target antimicrobial range of Zoocin A, CbpD-ves, and ScrM against (A) streptococcal species and (B) non-streptococcal species. The enzymes were added to cultures at OD_550_ ~ 0.2 at a final concentration of 30 µg/mL, and the OD_550_ of the treated cultures was compared with that of an untreated control 1 hour after enzyme addition (relative OD_550_).

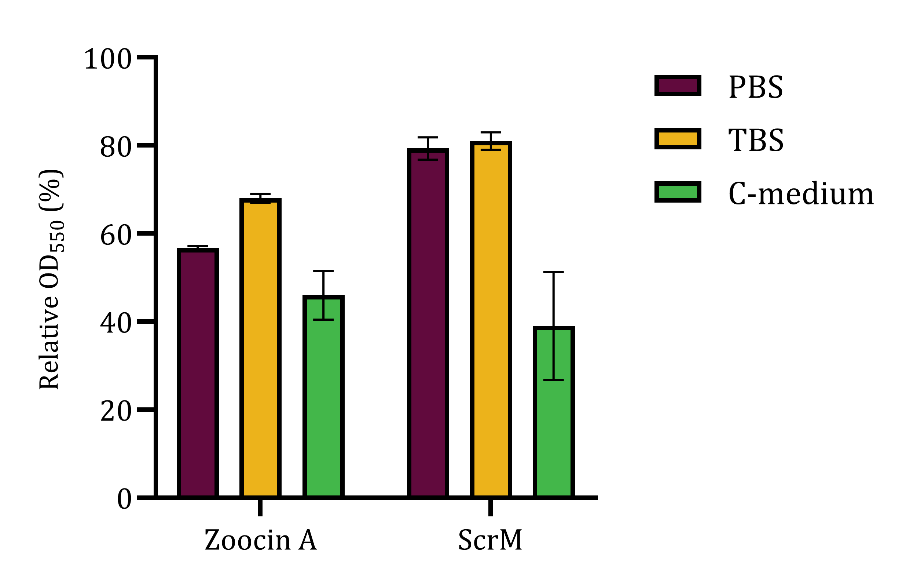

**Fig. S3** Activity of Zoocin A and ScrM in PBS, TBS, and C-medium. Cells were washed and suspended in the indicated solution prior to enzyme addition to a final concentration of 2 µg/mL. The relative OD_550_ was determined by comparing the OD_550_ measured 1 hour after treatment with that measured immediately before treatment.

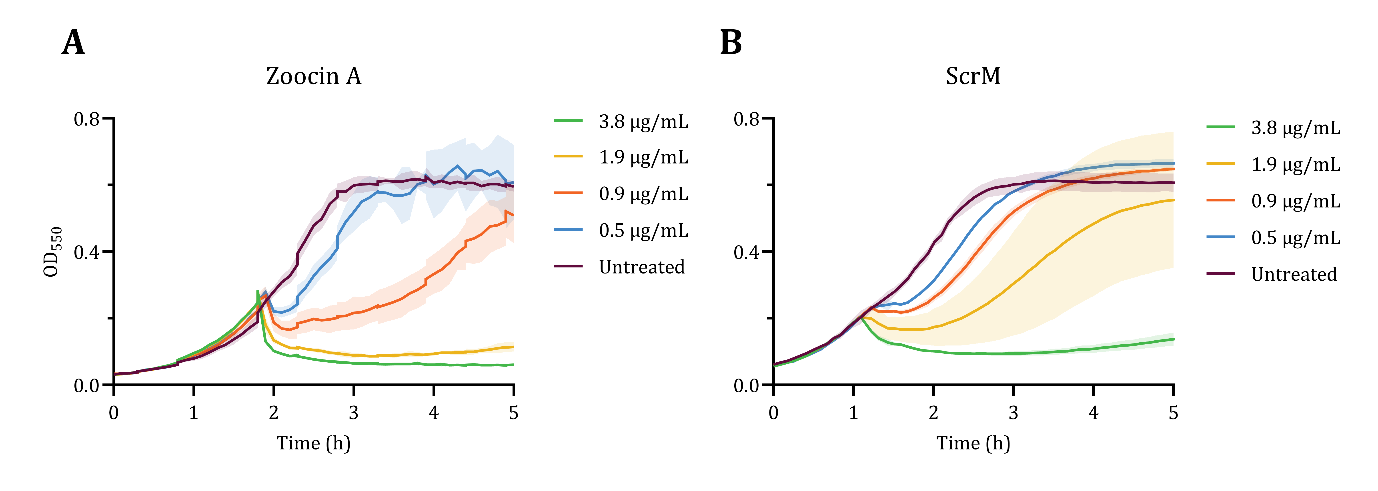

**Fig. S4** MIC₅₀ values of *S. agalactiae* for Zoocin A (A) and *S. dysgalactiae* for ScrM (B). Two-fold dilution series of enzymes were added to actively growing cultures at OD_550_ = 0.2, resulting in final enzyme concentrations ranging from 30 – 0.06 µg/mL. An untreated culture was included as a control. The MIC₅₀ was determined to be approximately 1 µg/mL (0.9 µg/mL) for Zoocin A against *S. agalactiae* and 2 µg/mL (1.9 µg/mL) for ScrM against *S. dysgalactiae*.

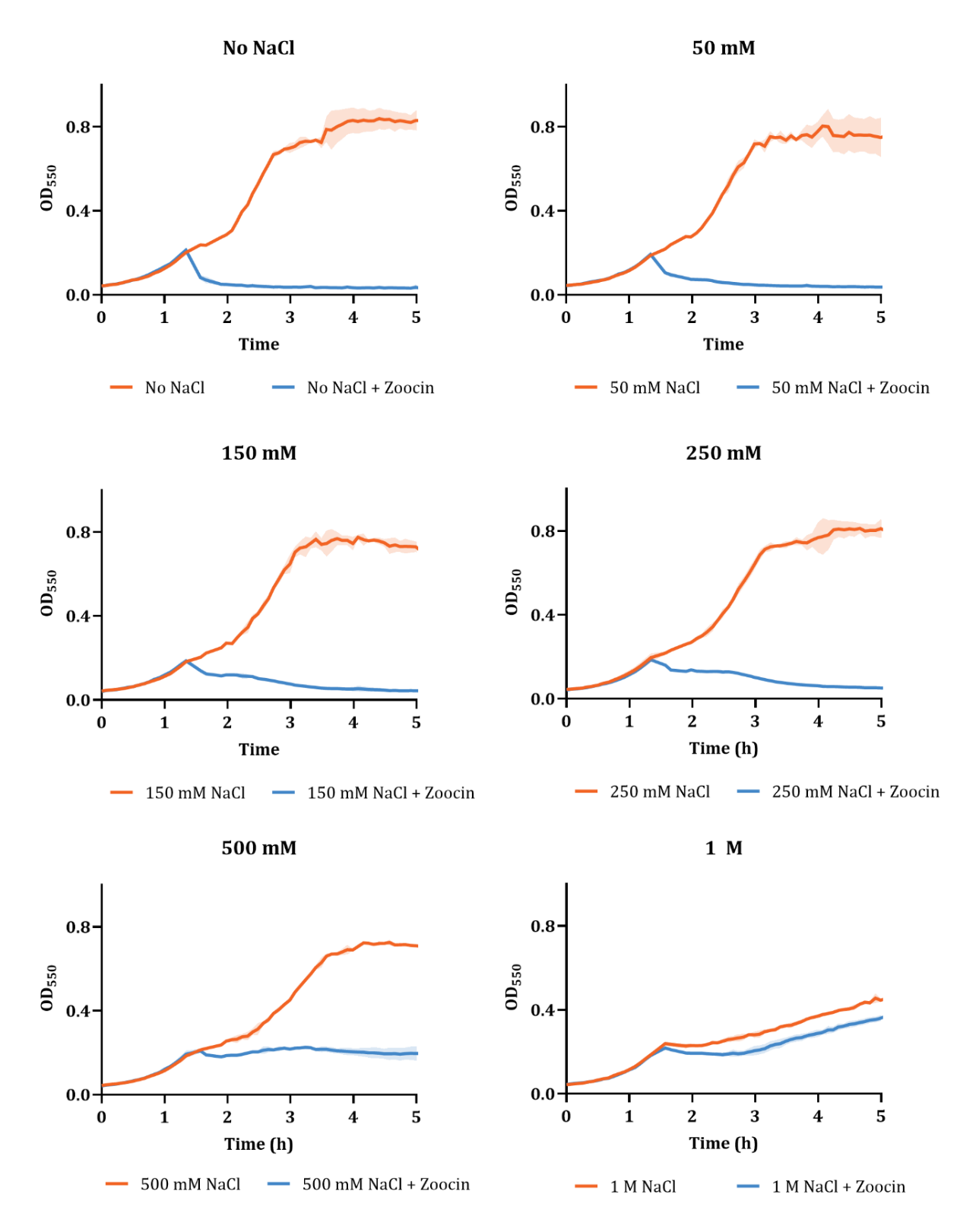

**Fig. S5** Effect of NaCl concentration on Zoocin A activity. Growth curves of cultures treated with Zoocin A at OD_550_ = 0.2, to a final concentration of 2 µg/mL, in the presence of 0, 50, 150, 250, 500, and 1000 mM NaCl. For each NaCl concentration, a corresponding enzyme-free control containing the same concentration of NaCl was included.

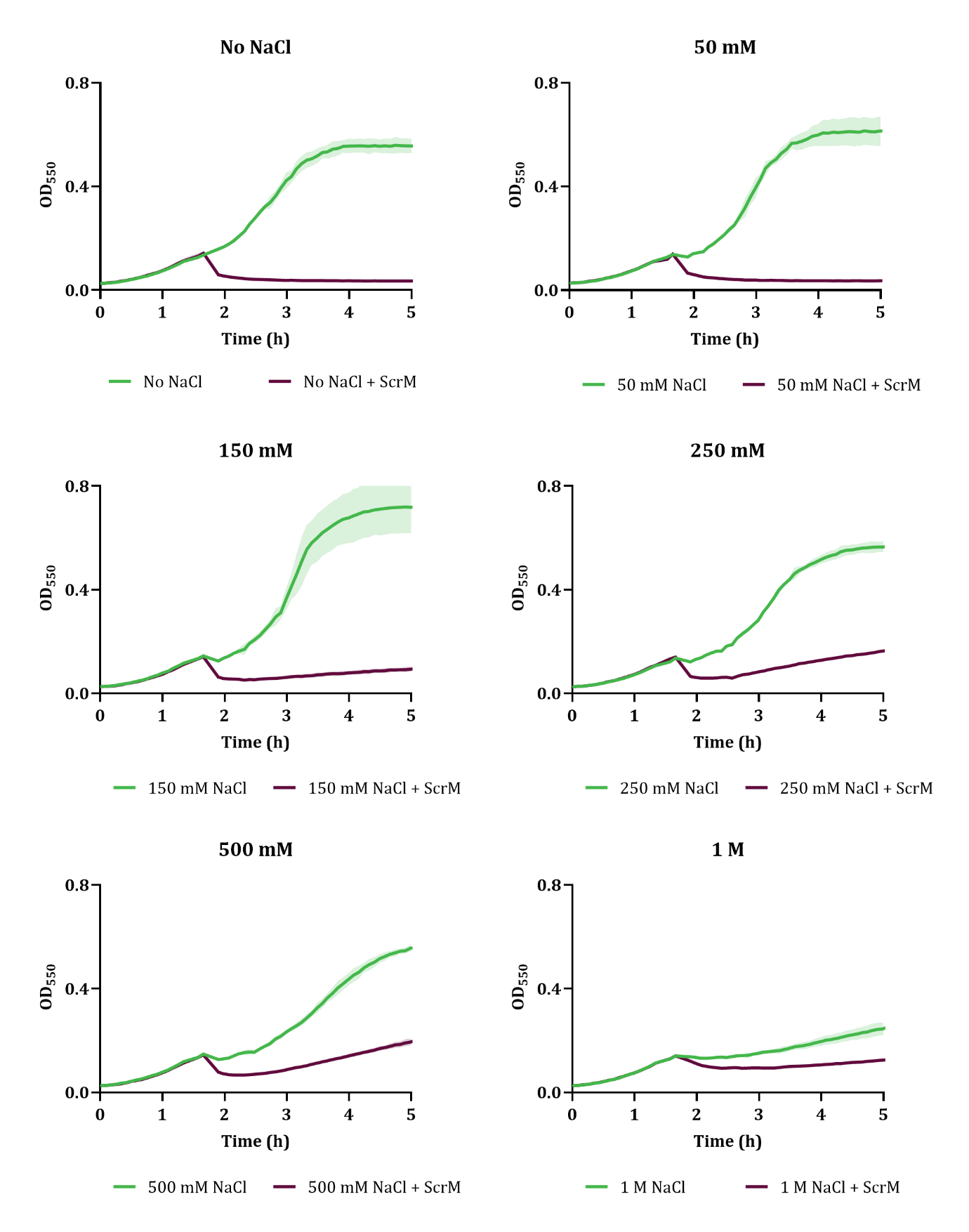

**Fig. S6** Effect of NaCl concentration on ScrM activity. Growth curves of cultures treated with ScrM at OD_550_ = 0.2, to a final concentration of 4 µg/mL, in the presence of 0, 50, 150, 250, 500, and 1000 mM NaCl. For each NaCl concentration, a corresponding enzyme-free control containing the same concentration of NaCl was included.

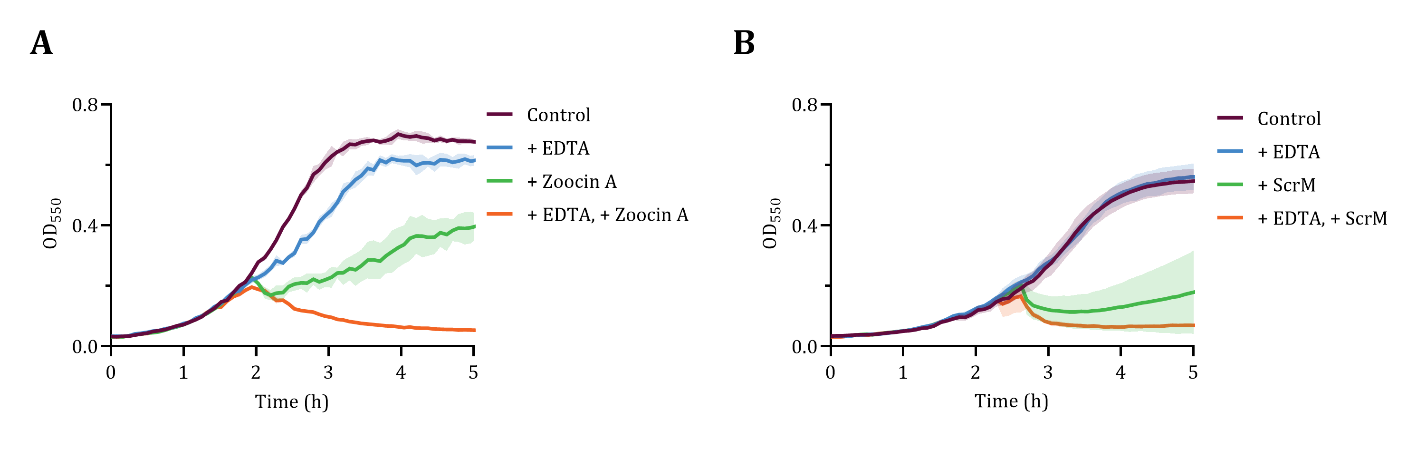

**Fig. S7** Effect of EDTA on the activity of Zoocin A (A) and ScrM (B). The chelating agent ethylenediaminetetraacetic (EDTA) was added to the cultures and enzyme stocks to a final concentration of 1 mM 30 minutes before adding the enzymes to the cultures (at OD_550_ = 0.2). Zoocin A and ScrM were added at MIC_50_ concentrations (1 µg/mL and 2 µg/mL, respectively). Controls included untreated cultures (Control), cultures treated with EDTA alone (+ EDTA), and cultures treated with the enzymes alone (+ Zoocin A/ScrM).

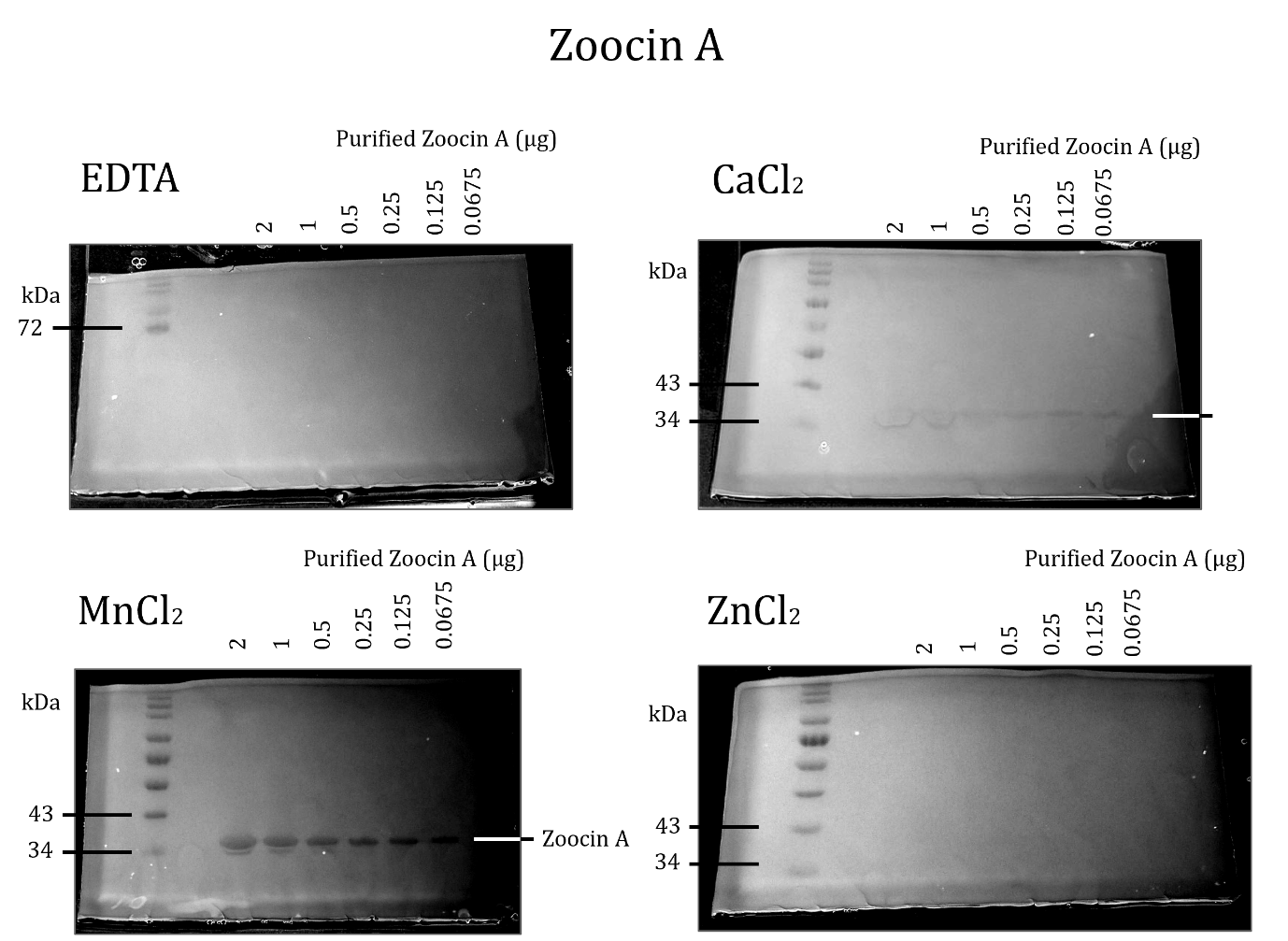

**Fig. S8** Full-size zymography gels showing the effect of divalent cations on Zoocin A activity. The full-size gel corresponding to Figure 5 (A) is shown. The gel was treated with 1 mM EDTA to remove divalent cations and then submerged in buffers containing 1 mM CaCl₂, MnCl₂, or ZnCl₂. A gel maintained in EDTA was included as a control. Molecular weights (kDa) are indicated on the left, and the amount of Zoocin A loaded is indicated at the top of each gel.

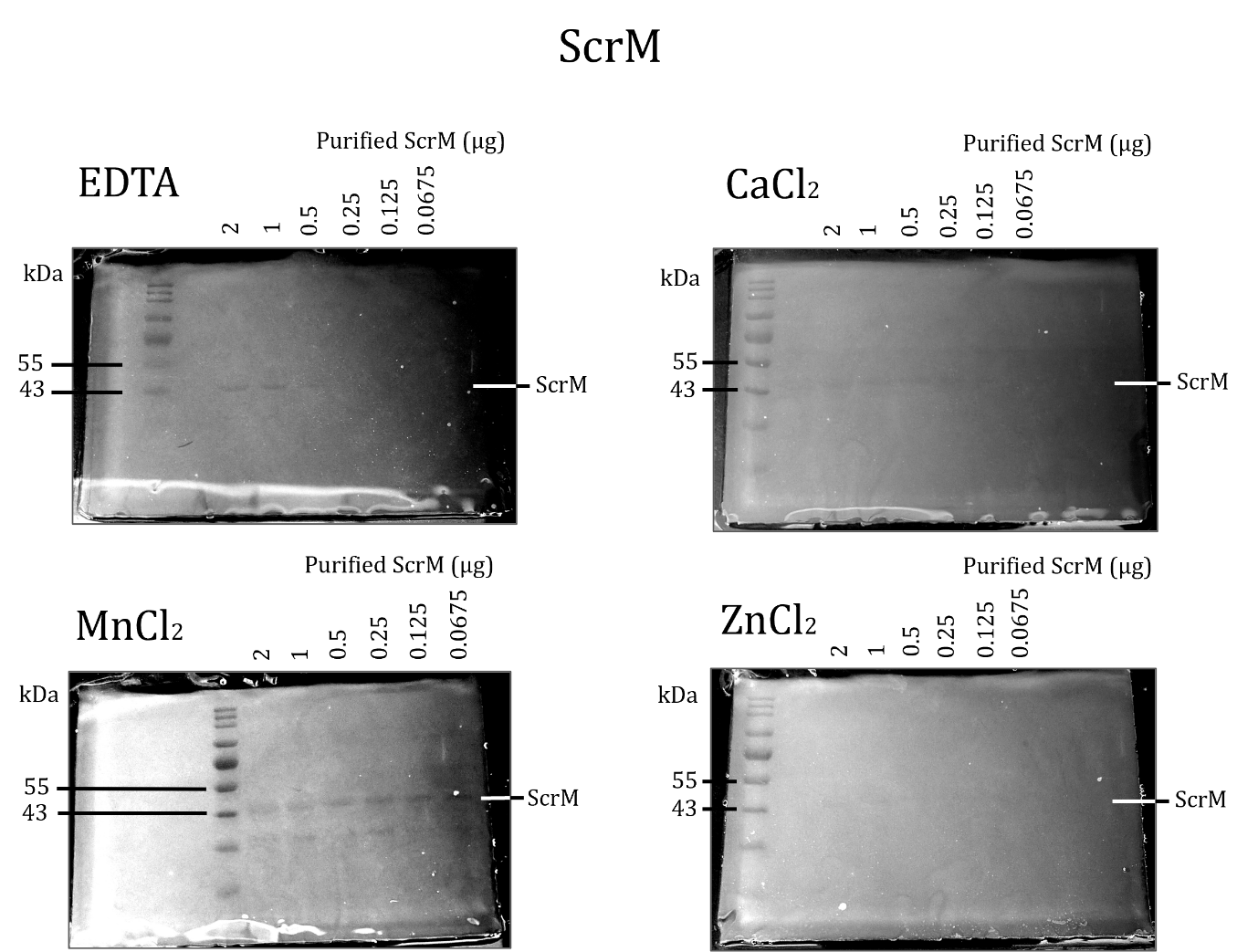

**Fig. S9** Full-size zymography gels showing the effect of divalent cations on ScrM activity. The full-size gel corresponding to Figure 5 (B) is shown. The gel was treated with 1 mM EDTA to remove divalent cations and then submerged in buffers containing 1mM CaCl₂, MnCl₂, or ZnCl₂. A gel maintained in EDTA was included as a control. Molecular weights (kDa) are indicated on the left, and the amount of ScrM loaded is indicated at the top of each gel.

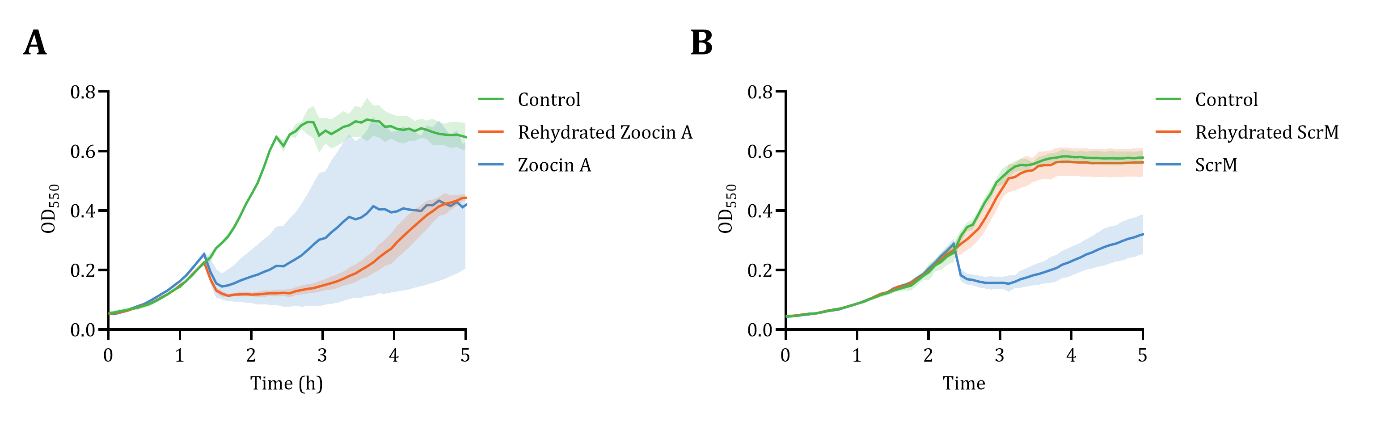

**Fig. S10** Effect of drying on the activity of Zoocin A (A) and ScrM (B). The enzymes were vacuum-dried for approximately 1 hour before being resuspended in dH_2_O and added to cultures at OD₅₅₀ = 0.2 to final concentrations of 1 µg/mL for Zoocin A and 2 µg/mL for ScrM. Controls included an untreated culture and a culture treated with the corresponding concentration of undried enzyme.
