## Supplemental Sequences for "Exploring the antimicrobial potential of streptococcal fratricins"

**Amino acid FASTA sequences of putative fratricins used in domain organisation analysis (Fig. 1)**

>WP_125375772.1 GBS Bsp-like repeat-containing protein [Streptococcus intermedius]

MKKYSQLFLLSSALLSVFSAQLTVQASEANNASKSSNPTTLASDVTVNVSGNSVSINYIRSQAQNPYTVSHAVWSDENGQDDLKWYSASQTSTTVDLSQHSGYGTFHVHTYININGKMIGLNGTTFTLNRPTSQTKITTAGQIGQIHFTRNKDQKNSKIVHAVWSEENGSDDINWYDAEQEITKFDFSKHKGYGRYFVDTYENKNGKMIYQSGTTFNLEKPNPTIQTSFPEPGIMDIRIKNIPETIYKLTVPTWSDKNGQDDLQWYAATKNPDGSYNVRVELKKHNYDTGTYHIHVYGESYVKPEFTGLAGTTVQVRSDQLPSEEDQKPLFSVENINQEQGTYTVKIKETSKSKSIQSVRVPIWSTTNQSNIKWYTATDNGDGTFSTTFNIRNHQALSGTYINHIYVKYKDGSEHSYATDSVSMSAEQIKTKVAVTKRSSYNYEVTVTDAYGDGNIILPTWSEVNGQDDIQWYTATKTGNGIYKFTIDTQKHTGSGLFHTHVYRNLNGKMIGLTGTSYQVEKPVISFQPNYAAATTYPKGQCTWGAKALAPWAGNYWGNGGDWATSARRAGFKTGTTPQVGAIICWTDGSYGHVGVVTHVASNTRIQIQEANYAGKQYIGNFRGWFNPHAAGQGVVSYIYPK

>WP_020997517.1 GBS Bsp-like repeat-containing protein [Streptococcus constellatus]

MKKYSQLFLLSSALLSVFSAQLTVQASEANNASKNSNPTTLASDVTVNVSGNSVSINYIRSQAQNPYTISHAVWSDENGQDDLKWYSASQTSTTVDLSQHSGYGTFHVHTYININGKMIGLNGTTFKVEKPANSTVSTVFGKNAQISFTRNKDQTNANILHAVWSDEKGQDDIKWYTAGQHKTEIDLSNHRGYGIYHVHTYENKNGKMTGLNGTTFNLEKPNPTIQTSFPEPGIMDIRIKNVPETIYKLTVPTWSDKNGQDDLQWYAATKNPDGSYNVRVELKKHNYDTGTYHIHLYGESYVKPEFTGLAGTTVQVRSDQLPSEEEQKPFFSVENINQEQGTYTVKVSETSKSKPIQSVRVPIWSTSNQSNIKWYAATNNGDGTFTATFDIRNHQVLSGTYTNHIYVKYKDGSEHSYATDSVTMSAEKIKTKVSVNKRSTYLYEVTVTDAYGDGTITLPTWSEVNGQDDIQWYTATKAGNGIYKFTVDTQKHTGSGLFHTHVYRNLNGQMIGLAGTSYQVEKPAVSFQPNYAAATTYPKGQCTWGAKALAPWAGNYWGNGGDWAASARRAGFKTGTTPQVGAIICWTDGSYGHVGVVTHVASNTRIQIQEANYAGKQYIGNFRGWFNPHAAGQGVVSYIYPK

>WP_195323494.1 GBS Bsp-like repeat-containing protein [Streptococcus anginosus]

MKKYSQLFLLSSTILGVFSAQLTAQASEADNAQKSSNPATLASNITVNVSGNTASINYTRSQTQVPYTIYHAVWSDENGQDDIKWYSAPQTPTTAIDLRQHTGYGTFHVHTYININGKMVGLNGTTFKVEKPASGTVSTTVLGKTAQISFTRNKDQTNANILHAVWSDEKGQDDIKWYTAGQDTTEIDLSNHKGYGVYHVHTYESKNGKMIGLNGTTFNLEKPNPTIQTSFPEPGIMDIQIKNVPETIYKLTVPAWSDKNGQDDLQWYAATKNPDGSYNVRVELKKHNYDTGTYHIHLYGESYVKPEFTGLAGITAQVDADKLPSEEEQKPFFSVENINQEQGTYTVKVSETSKSKPIQSVRVPIWSTSNQSNIKWYAATNHGDGTFTAAFDIRNHQVLSGTYTNHIYVKYKDGSEHSYATDSVAMSAEKIKTKVSVNKRSTYLYEVTVTDAYGDGSITLPTWSEVNGQDDIQWYTATKAGNGIYKFTIDTQKHTGSGLFHTHVYRNLNGQMIGLTGTSYQVEKPAVSFQPNYAAATTYPKGQCTWGAKALAPWAGNYWGNGGDWAASARRAGFKTGTTPQVGAIICWTDGGYGHVGVVTHVASNTRIQIQEANYAGKQYIGNFRGWFNPHAAGQGAVSYIYPK

>WP_214257845.1 GBS Bsp-like repeat-containing protein [Streptococcus lutetiensis]

MNKIKSALLSTAVLSMAFLTNSAMADQQSDALVTKTPFDVSSKTFEVTAQQSVNGKAIKSIDVAVWSEENGQDDIKWYSSSNVSNGQVKVQFNLANHGNHAGNYNTHVYTNYTDGSRSGVVMENTLISPKAPTLSLKDGGIRMTTDIYAPTNGTLYYAVWSEENDQDDIKWYADNGTGTTIADLKNHSGYGKYSIHTYVSQNGKMTGISCQDITINKQEISYQINKVNDTTYDVLVTNVPEYITSITVPVWSNANGQDDIKWYTTTKIGEGTYKAQINLGNHGYISGDYSVHIYGHNAISNNFEGLRVTSGFNAVNVPKPNATPSPRITTYINETNTYPVGQCTWAVKSLAPWIPNWLGNAGGWAVNARAKGFRVGTTPRVGSIVVWPHDGNGYGHVAYVTDVSSNTRIQVKESNYAGKQYIANFRGWFNPLDSFWGGDVSYIYPD

>WP_058621559.1 GBS Bsp-like repeat-containing protein [Streptococcus gallolyticus]

MSRLKTALLSTVVLSMAFLTNSVMADQQGDALVTKTPFDVASKTFEVTAQQSDNGKAIKGINVAIWSEENGQDDIKWYSTSDISNGQARVRFNLANHGNHAGNYITHVYTNYADGSRSGVVMENTWISPKAPTLSLKDGGIRMSTDIYAPSNGQIKYAVWSEENDQDDLKWYEDSGKGITQADLKNHSGYGRYFVHTYLSQGGKMIGINGQDIVINKQDVSYQINKINDTTYDVLVTNVPEYITSITVPVWSNVNGQDDIKWYTTTKIGEGTYKAQVNLANHGYTSGDYSVHIYGQNAINNNFEGLKGTSGFRADNVPNQNTTPTPRITTYINETNTYPVGQCTWAVKTLAPWIPNWLGNAGGWATNAKTKGFRTGSTPRVGAIAVWPNDGGGYGHVAYVADVASNTRIQVKESNYAGNQYIANFRGWFNPLDSFWGGSVTYIYPD

>WP_074567125.1 GBS Bsp-like repeat-containing protein [Streptococcus equinus]

MNKIKSALLSTAVLSMAILSNSVMADQQNDALVTKTPFDAVSKTFEVTAQQSPNGKAIKSIDVAVWSEENGQDDLKWYSSSDVSNGQVKVQFNLANHGNHAGNYNTHVYTNYSDGSRSGVVMENTLISPKAPTLSLKDGGIRMTTDIYAPTNGTLYYAVWSEENDQDDIKWYADNGTGATIADLKNHSGYGKYNVHTYVSQDGKMTGISCQDITINQQEISYQINKVNATTYDVLVTNVPEYITSITVPVWSNANGQDDIKWYTTTKIGEGTYKAQINLANHGYASGDYSVHIYGQNAISNSFEGLKGTSGFRAENVPNQNTAPTPRITTYINETNTYPVGQCTWGVKSLAPWIPNWLGNAGGWAVNARAKGFRVGTTPRVGSIVVWPHDGGGYGHVAYVTHVSSNTRIQVKEANYAGKHYIGNFRGWFNPLDSFWGGDVSYIYPD

>WP_093528547.1 GBS Bsp-like repeat-containing protein [Streptococcus gallolyticus subsp. macedonicus]

MSRLKTALLSTVVLGATCLSHPVFADQQGNAIVSKTNFDSASKTFEVTAQQSDNGKAIKGINVAIWSEENGQDDLKWYTNSNVSGGQVKVQFNLANHGNRAGNYITHVYTNYTDGSCSGVALENTKINPKALKLSVKNGAIQMTTDIYAPSNGQIKYAVWSEENDQDDLKWYEDSGKGITQADLKNHSGYGRYFVHTYLSQGGKMIGINGQDIVINKQDVSYQINKINDTTYDVLVTNVPEYITSITVPIWSNANGQDDIKWYTTTKIGEGTYKAQVNLANHGYASGDYSVHIYGQNAINNNFEGLKGTSGFRADNVPNQNTTPTPRITTYINETNTYPVGQCTWAVKSLAPWIPNWLGNAGGWATNAKAKGFRTGSTPRVGAIAVWPNDGGGYGHVAYVTDAASSTRIQVKESNYAGNQYIANFRGWFNPLDSFWGGSVTYIYPD

>WP_248497737.1 GBS Bsp-like repeat-containing protein [Streptococcus gallolyticus subsp. pasteurianus]

MSKLKTALLSTVVLGATCLSHPVFADQQGNAIVSKTNFDSASKTFEVTAQQSDNGKAIKGINVAIWSEENGQDDLKWYTNSNVSGGQVKVQFNLANHGNRAGNYITHVYTNYTDGSCSGVALENTKINPKAPQLSVKNGAIQMTTDIYAPSNGQIKYAVWSEENDQDDLRWYEDSGKGITQADLKNHSGYGRYFVHTYLSQGGKMIGINGQDIVINKQEVSYQINKVNDTTYDVLVTNVPEYITSIAVPTWSNANGQDDIKWYTTTKVDNGTYKAQINLANHGYASGDYSVHIYGQNAITNNFEGLKGTSGFRADNVPNQNATPTPRITTYINETNTYPVGQCTWAVKSLAPWIPNWLGNAGGWATNAKAKGFRTGSTPRVGSIAVWPNDGGGYGHVAYVTDVASNTRIQVKESNYAGNQYIANFRGWFNPLDSFWGGSVTYIYPD

>WP_077497030.1 GBS Bsp-like repeat-containing protein [Streptococcus gallolyticus subsp. gallolyticus]

MSKLKTALLSTVVLGATCLSHPIFADQQGNAIVSKTNFDSASKTFEVTAQQSDNGKAIKGINVAIWSEENGQDDLKWYTNSNVSGGQVKVQFNLANHGNRAGNYITHVYTNYTDGSCSGVALENTKINPKAPQLSVKNGAIQMTTDIYAPSNGQIKYAVWSEENDQDDLRWYEDSGKGITQADLKNHSGYGRYFVHTYLSQGGKMIGINGQDIVINKQEVSYQINKVNDTTYDVLVTNVPEYITSIAVPTWSNANGQDDIKWYTTTKVDNGTYKAKINLANHGYASGDYSVHIYGQNAITNNFEGLKGTSGFRADNVPNQNTTPTPRITTYINETNTYPVGQCTWAVKSLAPWIPNWLGNAGGWATNAKAKGFRTGSTPRVGSIAVWPNDGGGYGHVAYVTDVASNTRIQVKESNYAGNQYIANFRGWFNPLDSFWGGSVTYIYPD

>MCO4485798.1 glucan binding protein [Streptococcus infantarius]

MNKIKSALLSTAVLGMAFLTNSAMADQQSDALVTKTPFDVSSKTFEVTAQQSVNGKAIKSIDVAVWSEENGQDDIKWYSSSNVSNGQVKVQFNLANHGNHAGNYNTHVYTNYTDGSRSGVVMENTLISPKAPTLSLKDGGIRMTTDIYAPTNGTLYYAVWSEENDQDDIKWYADNGTGTTIADLKNHSGYGKYSIHTYVSQNGKMTGISCQDITINKQEISYQINKVNDTTYDVLVTNVPEYITSITVPVWSNANGQDDIKWYTTTKIGEGTYKAQINLANHGYASGDYSVHIYGQNAISNNFEGLRVTSGFNAVNVPKPNATPSPRITTYINETNTYPVGQCTWAVKSLAPWIPNWLGNAGGWAVNARAKGFRVGTTPRVGSIVVWPHDGNGYGHVAYVTDVSSNTRIQVKESNYAGKQYIANFRGWFNPLDSFWGGDVSYIYPD

>WP_074567125.1 GBS Bsp-like repeat-containing protein [Streptococcus equinus]

MNKIKSALLSTAVLSMAILSNSVMADQQNDALVTKTPFDAVSKTFEVTAQQSPNGKAIKSIDVAVWSEENGQDDLKWYSSSDVSNGQVKVQFNLANHGNHAGNYNTHVYTNYSDGSRSGVVMENTLISPKAPTLSLKDGGIRMTTDIYAPTNGTLYYAVWSEENDQDDIKWYADNGTGATIADLKNHSGYGKYNVHTYVSQDGKMTGISCQDITINQQEISYQINKVNATTYDVLVTNVPEYITSITVPVWSNANGQDDIKWYTTTKIGEGTYKAQINLANHGYASGDYSVHIYGQNAISNSFEGLKGTSGFRAENVPNQNTAPTPRITTYINETNTYPVGQCTWGVKSLAPWIPNWLGNAGGWAVNARAKGFRVGTTPRVGSIVVWPHDGGGYGHVAYVTHVSSNTRIQVKEANYAGKHYIGNFRGWFNPLDSFWGGDVSYIYPD

>tr|Q9KGZ2|Q9KGZ2_STREE Choline binding protein D OS=Streptococcus pneumoniae OX=1313 GN=cbpD PE=4 SV=1

MKILPFIARGTSYYLKMSVKKLVPFLVVGLMLAAGDSVYAYSRGNGSIARGDDYPAYYKNGSQEIDQWRMYSRQCTSFVAFRLSNVNGFEIPAAYGNANEWGHRARREGYRVDNTPTIGSITWSTAGTYGHVAWVSNVMGDQIEIEEYNYGYTESYNKRVIKANTMTGFIHFKDLDGGSVGNSQSSTSTGGTHYFKTKSAIKTEPLASGTVIDYYYPGEKVHYDQILEKDGYKWLSYTAYNGSYRYVQLEAVNKNPLGNSVLSSTGGTHYFKTKSAIKTEPLVSATVIDYYYPGEKVHYDQILEKDGYKWLSYTAYNGSRRYIQLEGVTSSQNYQNQSGNISSYGSHSSSTVGWKKINGSWYHFKSNGSKSTGWLKDGSSWYYLKLSGEMQTGWLKENGLWYYLGSSGAMKTGWYQVSGKWYYSYSSGALAVNTTVDGYRVNSDGERV

>WP_000866454.1 choline binding-anchored murein hydrolase CbpD [Streptococcus mitis]

MKVLPFKVTETGFSLRKSVKKVVPFLVVGLMLAASDSVYAYSGGNGSIARGDDYPAYYKNGSQEIDQWRMYSRQCTSFAAFRLSNVNGFEIPRAYGNANEWGHRARREGYRVDNTPTIGSIAWSTAGTYGHVAWVSNVMGDQIEIEEYNYGYTEAYNKRIIKANTMTGFIHFKDLAGGSVGNSQTSASTGGTHYFKSKAAIKNQPLASATAIDYYYPGEKVHYDQILEKDGYKWLSYTAYNGSRRYIQLEGVTSSQNYQNQSGNISSYGSNSSSTVGWKKINGSWYHFKSNGSKSTGWLKDGSSWYYLKSSGEMQTGWLKENGSWYYLDSSGAMKTGWYQVSGKWYYSYSSGVLAVNTTVDGYRVNSDGERV

>WP_000703688.1 SH3 domain-containing protein [Streptococcus oralis]

MKISPFTVTETGVSFRKSVKKVVPFLAVGLMLVAGDSVYAYSGGNGSFARGDDYPLHYKNGSVEIDQWRMYSRQCTSFVAFRLSSVNGFEIPPGYGNANEWGYRARREGYRVDSKPEVGSIAWSTEDYYGHVAWVSNVMG

DQIEIEEYNYGVRERYNRRIVKASSMTGFIHFKDLVGNDGRTGSPIESGLASSGTHTFTQKSAIRNQPSSTAQVIDYYYPGENVSYDQIVEKDGYKWLSYLSYSGSRRYVQYTETESVENGWRKQNGIWNYLENGKLATGWKKINGSWYHFKDNGTMSTGWVKDGSYWYYLKASGEMQTGWLNENGTWYYLESSGAMKSSQWFQVGGKYYYVNASGALAVNTTVDGYRVDSNGARI

>WP_011836292.1 murein hydrolase LytF [Streptococcus sanguinis]

MKKYQKLFLLSGAVLGLFASHSTVHATETPESEVKILGPTDRASNVDVTVNDRTVSISYTRSQAQEPYLIAHAVWSEENGQDDLKWYDTPQTPITDIDLSNHPGYGTFHVHTYIRIGERLVALNATTFHVDKPASKTNASVSGGTGRISFSRNKDQTHSKIVHAVWSEEKGSDDLQWYDAGEESTEFNLSKHRGYGRYFVDTYENVNGKMIFLSGTTFHLENPKPIIETSFPQTGIMEITVKNVPDTMYKVVLPTWSDKKGQDDLQWYEASKNPDGSYSARVELRKHNYDTGTYNIHLYGQSYVQPESTGITGTTVQVDSGKLPSDEEQKPLFTVENINPKQGTYTVKTTETSLSKPIQSVRVPIWSTSNQSNLKWYAATPNGDGSFSATFDIRNHQALSGTYNNHVYVTYKDGSERSYAADAVSMSTDQIQAKVAVRKTEANRYEVTVTDAYGDGDIILPTWSEVNGQDDIKWYTANKVGNGTYKFTVDTQNHKGSGLFNTHVYRRKAGQLIGLTGTSYQVEKSSVQSANIQPNYAAANATTYPVGQCTWGAKALAPWAGNYWGNGGQWAASARRAGFRTGSTPEVGAIACWDDGGYGHVGVVTHVESNTRIQIQESNYLGKQYISNFRGWFDPTASYWGRLTYIYPK

>WP_013904392.1 GBS Bsp-like repeat-containing protein [Streptococcus parasanguinis]

MQTIKKFVVLTCATWGMQASIAHADQATNAIYAEKGQLMIHLGNVPDKYQKIQVPVWSDQNGQDDLIWYPVERSDKGFDLQVPLTNHSDQAGLYHVHVYGVEANEQLTGLYPLTTSVQQKDLASSQPKITITPISSQEFDIQLKLFEDVDEIVFPIWSEENGQDDLVWYPAKKIAPGHFQLRFQVQKHKGSGLFHLHVYQKSQGQLKGLFPTSFQVEKAKPRLTPLATHPGNTYPIGECTWGAKELAPWAHNWWGNGGMWAASARAAGFRTGDTPEVGAIACWDNGGYGHVALVTDVEHDQKIQIQEANYNGHRYIDNFRGWFDPTNPIWGTVTYIYPD

>MBF1698418.1 GBS Bsp-like repeat-containing protein [Streptococcus cristatus]

MKKYQKLFLLSGAVLGLFASHSTVHATETPESEVKILGPTDRASNVDVTVNDRTASISYTRSQAQEPYLIAHAVWSEENGQDDLKWYDTPQTPITDIDLSNHPGYGTFHVHTYIRIGERLVALNATTFHVDKPASKTNASVSGGTGRISFSRNKDQTHSKIVHAVWSEEKGSDDLQWYDAGEESTEFNLSKHRGYGRYFVDTYENVNGKMIFLSGTTFHLENPKPIIETSFPQTGIMEITVKNVPDTMHRIVLPTWSDKKGQDDLQWYEASKNSDGSYSARVELRKHNYDTGTYNIHLYGQSYVQPESTGITGTTVQVDSGKLPSDEEQKPLFTVENINPKQGTYTVKTTETSLSKPIQSVRVPIWSTSNQSNLKWYAATPNGDGSFSASFDIRNHQALSGTYNNHVYVTYKDGSERSYAADAVSMSTDQIQAKVAVRKTEANRYEVTVTDAYGDGDIILPTWSEVNGQDDIKWYTANKVGNGTYKFTVDTQKHKGSGLFHTHVYRRKAGQLTGLTGTSYQVEKSSVQSANIQPNYAAANATTYPVGQCTWGAKALAPWAGNYWGNGGQWAASARRAGFRTGSTPEVGAIACWDDGGYGHVGVVTHVESNTRIQIQESNYLGKQYISNFRGWFDPTASYWGRLTYIYPK

>WP_223334409.1 SH3 domain-containing protein [Streptococcus infantis]

MTYSKKGTFLKRTTVGTASLLLAGAFLFGGGAVHANQANNGSLAQGDDYPYYYKNGSQEIDKWRMYSRQCTSFAAFRLSSVNGFELPAAYGNANQWGYRAKREGYRVDQHPAIGSIAWDASGQYGHVAWVSNVLGDMIEIEEYNYGIREKYNRRIVKASSMTGFIHFKDVGQASTGNHQATLVSSGTHVFTEKAAIMNQPSQAAEIIDYYYPGESVHYDQTLEKDGYKWISYLSRTGNRRYVQIAKTATAKNGWEKQGTIWYYYENDKVALGWKKINGSWYYFQTSGAMRTGWLKDGSKWYYLKSSGEMQTGWLKDKGIWYYLESSGAMKASQWFQVSGKYYYVNASGELAVDTTIDGFQVDSNGARVEESPQ

>WP_012131057.1 GBS Bsp-like repeat-containing protein [Streptococcus gordonii]

MKKYKNFLLLSATLLTSLSAPLAAANEQFSPSQLTAMTSEVYVQLEGQTAIVQYQPTAAQEAYTIQHAVWSEEGGQDDIVWYSADQSTTRIDLSQHKGQGKFLIETYLNVNSVKYHLSSKSVFLNKPVQSTEKTVPEQQDSTSSNNATPQAPAEPKPTNPAPTPAPAPATNRVENPRNSYPKVEERNKSLPPQKVAQPKVAPPQAEPNNNKPLQASAARVPAEPVRDSKPLISITNINKENGTYTVRVQENPYSKKIKSVQVPIWSTANQSNLKWYEASKIGNGSFSVQFDIRNHQQIYGNYINHVYITYEDGSRVGYVAENVNLPQPAPQPLTPSLQVVHKGQSTYQVILQNSFGDGHILFPIWSDINGQDDLRWYTANSLGNGRYSLDFNANNHIGDGLYHVHVYRQSGGKVTGVMASNFQVTRPTVAVRKPLNHTTDPSNTYPVGECTWGAKALAPWAGNWWGNGGDWAASARRAGFRVGSTPQVGAIACWTDGGYGHVGVVTHVDSHKRIQIQESNYAGQRYIANFRGWFDPTIAQGTVSYIYPN

>WP_006144633.1 SH3 domain-containing protein [Streptococcus peroris]

MTFSKKKTFLKRTTVGTASLLLAGTLLFGGGTVHANQGNNGSLARGDDYPYYYKNGNQEIDKWRLYSRQCTSFAAFRLSSVNGFELPAAYGHAKEWGYRAQREGYRVDQKPAIGSIAWSTRGSYGHVAWVSNVIGDMVEIEEYNYDYTETYHRRTVKASSMTGFIHFKDVGNTSVENYQSTPASSGTHYFTEKAAIMDQPSLTSTIIDYYEPGMSVHYDQTLEKDGYKWISYISRSGNRRYIPLTKTAISKNGWEKQGTTWYYYENDKVALGWKKIDGSWYYFQTSGAMRTGWLKDGSKWYYLKSSGEMQTGWLKDKGIWYYLESSGAMKSSQWFQVSGKYYYVNASGELAVDTTIDGFQVDSNGARVEKSTQ

>WP_223334409.1 SH3 domain-containing protein [Streptococcus infantis]

MTYSKKGTFLKRTTVGTASLLLAGAFLFGGGAVHANQANNGSLAQGDDYPYYYKNGSQEIDKWRMYSRQCTSFAAFRLSSVNGFELPAAYGNANQWGYRAKREGYRVDQHPAIGSIAWDASGQYGHVAWVSNVLGDMIEIEEYNYGIREKYNRRIVKASSMTGFIHFKDVGQASTGNHQATLVSSGTHVFTEKAAIMNQPSQAAEIIDYYYPGESVHYDQTLEKDGYKWISYLSRTGNRRYVQIAKTATAKNGWEKQGTIWYYYENDKVALGWKKINGSWYYFQTSGAMRTGWLKDGSKWYYLKSSGEMQTGWLKDKGIWYYLESSGAMKASQWFQVSGKYYYVNASGELAVDTTIDGFQVDSNGARVEESPQ

>WP_231911642.1 GBS Bsp-like repeat-containing protein [Streptococcus australis]

MKKLLLAACLFWAGASQVARAEQNVNYIYKEKGQLVVHLGTIPDRFQSVSVPIWSEQNGQDDLIWYPVQRGENGFDLQVPLTNHLDQAGLYHIHVYGIHSNGNIEGLFPLQTIVEKKDLASSQPKITVQPSGAQLFEIQLQLFEDVEEVLFPIWSEQDGQDDLIWYPAKRMAPGRYQLSFNAEKHKGNGTFHLHVYQKNKGHLKGLLATEFQVERIKPAPLVTQPDNYYPIGECTWAAKELAPWSQNWWGNGGMWTTSARAAGFRTGNTPEVGAIACWDNGGYGHVAVVTEVEHNQKIQVKEANYNGNRTINNFRGWFDPTNVVWGTVSYIYPN

>WP_217473448.1 GBS Bsp-like repeat-containing protein [Streptococcus sinensis]

MKKYQKLFLLSGAAIGLFASHSTAQAVASTEAGSKILGPTDRASNVIVNVTGDTAQIHYARSKAQVPYTIYHAVWSDENGQDDLKWYTTPQTSSTAIDLRQHAGYGTFHVHTYININGKMIGLNGTTFTLNKPTSKTSVTTAGQIGRINFTRNKDQGNSKIVHAVWSDENGGDDLNWYEAGQDSTEFNLSKHKGYGKYFVDTYENKNGKMIYQSGTTFHLEKPNPTIQTSFPESGIMEILIKNLPDTMYKVTVPTWSENKGQDDLQWYEASKNSDGSYRVRVELKKHNYDTGTYHIHLYGESYVKPEFTGLAGTTATIDAGKLPSPEEQKPLFSVENVNHEQGTYTVKTSETATSKPIQSVRVPIWSTHNQSNIKWYEASNNGDGTFTAQFNIRNHQALSGNYINHIYVKYKDGSEHSYATDTVTLSAENIKTKVSVNKISAYNYEVTVADAFGPGTISLPTWSEVNGQDDIKWYTANKVGNGLYKFTINTQQHAGSGLFHTHVYRTLNGQLTGLTGTSYQVDKPTTPEPTLYTPDYAGASSYPHGQCTWGAKVLAPWAGPYWGNGGQWAASARAAGFRTGSIPQVGAIICWTDGGYGHVGVVTHVESNTRIQIKESNYAGKQYIGNFRGWFNPYAAGQGAVSYIYPK

>WP_018374822.1 GBS Bsp-like repeat-containing protein [Streptococcus orisratti]

MKLIKSFLFTSCVLGMVALGQSASAEEIGQANVSKTEYNTSSRTFDVLVSRKAGGKAIKTVEVAIWSEENGQDDLRWYSSSAISNNQTKLQFNIANHGNRAGNYITHVYTTFTDGQRVGTNLGSVNIKTSLNPVQVVDGQVKISSTLAPPSNAKFQTAVWSDENGQDDIKWYNWDTTNLSVALSNHKGYGTYHVHTYLNQDNKMQGLSGQSFTLEKPAPAKISTQVTKISDKTVDISISGLPNYISSVKIPTWSEVNGQDDIKWYQAQKVGEGNFKVRVNMVDHGFDLGTYHAHFYGKNSQTGNEEGLAATSFTIDNISGLENPVINLNQTNSTQGLFTVSILETLMSKKVISLKVVVTSKTNVSKTKTYEVKTSNQRANISVDLKQLTTTADNFSVTAYAIYSDNSQAIFNLAEQKFTPSVVTTPQPSTTNSASPRITRYINETNTYPVGQCTWGVKTLAPWIPNYLGNAKEWTNGAKNLGFRTGTTPEVGAIAVWPYDGGGYGHVAYVTAVESSTRIQVKEANYAGNQSIANYRNWFNPTASWCGGGTVFYIYPN

>WP_002261986.1 GBS Bsp-like repeat-containing protein [Streptococcus mutans]

MRKLKVALFASSILGMLAVSSYTAADTEDNQVTISHYNEQAGTFDVNAVQAANGKTIQSIDVAIWSEENGQDDLKWYHASNDGSNQLTVHFNAENHGSKVGSYIAHAYITYTDGNRVGVNLGKRKLSLSAPQLSLKQGGLQLFSKLKPSAADQLFSAVWSDENGQDDLHWYTADADGNTLAGYANHKGYGTYHVHTYLKQNGKMIPISAQDIDIPKPKVKIQIDKINDTSYDVVVNNVPPYISSVAIPVWSEQNGQDDLKWYQATKVADGIFKTTVYLKTHRFELGNYQAHIYGDSQLSKKLDGLGETHFNVPSIINYEDPQVTIDHYNINKGTFDVTVAETDNSKAIQSISAAVWSDANQANLYWYEAKQLANGKAAITVDVQKHGNQTGSYNVHVYVHYNDGTTSGHVLANQQLNQIVHYQPSAVRITAYMNEKNTYPVGQCTWGVKELAPWIPNWLGNGGQWASTVAVKGFKIGTVPKVGAIACWSDGGYGHVAYVTHVESNNRIQVKEANYKNQQYISNFRGWFDPTTSYLGRLTYIYPD

>WP_029237162.1 RICIN domain-containing protein [Streptococcus sobrinus]

MTRNKRKMRKHELKKQAFLAGLTTMAVGAVALSSGHVQADEQVSSSNPSSDVSLVTGQEANQTISDVQVNSENQEQAQGTTTGSQAQNEKQGQAQATGEQVSQNSTDENQTSLESHEVSGTGVQAESETDPTQVTPIETEVNKEAGQENSTVKRASVVAVVERKTANTVSAARGDDYPAYLKNAAPDSVIDPWRLYNRECTSFTAWRLSSANGFTIPGAYGNGGQWGYRARREGYRVDNNPAIGSVGWLDDGSYGHVAWISNVMGDNVEIEEYNYGWNHSYHKRVAHKSAFTGYIHFKDLAVVSKPSDGRYDVESLAPATDRTLSDGDYHIALATNTDYVLDALGGQAKNGTNAQLYKDTNDSKQVFTVKYLGGGYYKIAQKVTGKVLDVTAGSKDNKANVQFWEDNGTEAQQWIIKQSSGNNTYEIISRRSGLAIDVANGKVSNNSNIQMFIPNKTDAQKFKFLAVDNDAKRTIADGDYHIASALGNGLALDVENESKNDGANIQVYPNLTNSKEVFTVKYLNNGYYSITHKYSGKMIDDSGNGAFNGNNISTITPNGSESQQWIIKASAINGYFEIISKHKNKLLDVEAGIAKPRTNVHLYARNNTKSQAWKFVPVKKD

>WP_003087899.1 GBS Bsp-like repeat-containing protein [Streptococcus ratti]

MKKVKSILFSTCLLGILACSQNVKADSGSANLFISNYSKFSGVFDVNALDPEGGKTIKDLRVAIWSEEGGQDDMQWYSAHNTGNDSLIVHFNPVNHASKAGNYAVHAYVTYTDGSKQGINLGKQKISLAEPQLGLAQNGIEIFSKLKPSGQGKLVSAVWSDENGQDDLKWYEADAAGNALADYDSHKGYGTYHIHTYLKKDGEMQPVAAQTLQLPRPNVSCQINKIKDTSYEAVISNVPPYMTSIYVPVWSEQSGQDDIQWYQAKKEADGTFKAKITLKKHYFDLGTYQLHLYGSGPLREWKLEGLESSAFNVEKIAAYEEPQLAVHHYNLKEGTFDVSAAETDDSKMIKSISLAVWSDTDQANLYWYESSKLSNGQASITADVQKHGNKSGSYNVHAYIHYTDGTSTGHILPNQQLNALKPSSLETPAPKIMAYISETNTYPIGQCTWGVKELAPWAPNWLGNGGQWAANARAKGFRTGNEARVGAIACWDDGGYGHVAYVTQVENNNRIQVKEANYQNQQYIENFRGWFDPTASYWGRLTYIYPD

>WP_040841467.1 GBS Bsp-like repeat-containing protein [Streptococcus macacae]

MKRLKIMLLTTAFLGVITAFRYAAADTQATITHYDKQAGTFDLVLPQVSDGQIIKSVDLAVWSEENGQDDLKWYSAQQKENGQLTVHFSTEQHGNMAGSYLVHAYITYANGKRIGVNLGKRDFSLSEPQLSLGPNGIQIASKMKPAGSEQLLWAVWSDQKGQDDIKWYLADDDGKAIAHYADHKGYGTYHVHTYLKKDGKMIPIAAQKLELPKPHIKSQINKINDTSYEITVSDVPSYITSVSVPVWTEQNGQDDLQWYQASKTADGIFKAVVYLKSHRFGLGRYQVHFYGDSLLNLDLEGLGSANFEVPSIDHYEAPQVTIGPYDNHKGTFDINVAETNHSKVIKTVKAAVWSEANQENIHWYESQQISDGKAAIRADIQKHGNKTGTYHVHTYITYTDESSSGHVLADQQLNAIDLTPAPSVRITAYINERNTYPVGQCTWGVKELAPWIPNWLGNGGQWADNARAKGFRTGNEAKVGAVACWDDGGYGHVAYVSHVDGNHRIQVKEANYKNQQFISNFRGWFDPTASYWGRLTYIYPD

>WP_018030002.1 GBS Bsp-like repeat-containing protein [Streptococcus ferus]

MKKVKSMLFATCLLGALACSQHAQADSGGASLSISNYNEFAGVFDVNALAQEGGKSIKEMRVAVWSEENGQDDLRWYSPAVYGNQANLHVDVSNHGNRAGSYITHAYVTYTDGSSEGKDLGKQKMSLSNPTVNVTEKTIQLSSKIKPAGNGQLAVAIWSEVGGQDDLKWYSLDANGNLTVPITNHKGDGNYHIHTYLNQDNQSIGVGGQVVQVTQPKLGSQIAKLSDTSYELTVTNVPGHISSVTIPVWSDSNGQDDIVWYQTEKVDVSTYKVRVPLKKHGFALGRYQAHIYGNNQATGKLEGLSTTGFDVTQIVNYENPVGTISNVNAAKGTFDVVISETDHSKRIKSVDVAIWSEANQANMVWYSSNKVSNGKVSITADIQKHGNKSGSYNSHAYVTFMDGSRAGYILPTQQMTQLNLNQAPAPKITAYINEANTYPVGQCTWGVKQLAPWIPNWLGNGGQWTANARAKGFRTGSTPKVGAVVSWDDGGYGHVAYVTHVESNNRIQVKEANYKNQQYIANFRGWFDPTAPYLGRVSYIYPD

>MSU87553.1 CHAP domain-containing protein [Streptococcus dysgalactiae subsp. dysgalactiae]

MKKIHQLLVSGAILLSVNGAVSSVASTLNADHTGVVHAAVLGDNYPSKWKKGSGIDSWNMYVRQCTSFVAFRLSSANGFQLPKGYGNACTWGHIAKKQGYTVNKTPKVGAVAWFDTNAFQSHATYGHVAWVAEVRGDSVVIEEYNYNAGQGPEKYHKRQIPKNHVSGYIHFKDLPSGEASKSQTKEQQVSKEAVKKGGTYHFTERTPVKAQAQLTSPDLAYYNPGQSVHYDQAMTVDGHEWISYLSFSGSRRYIPIKKAEQKTQQVSETTSPINIGDRVTFPGVFRVDRIVNNLLVSEELAGGGATSLNWIDPSPLDETDRKGVKAGNQILQAGEFFVIPGNYRVLKVDRPSNGIYVKIGSRGTWLTADKASKLP

>KKC19126.1 amidase [Streptococcus dysgalactiae subsp. equisimilis]

MKKIHQLLVSGAILLSVNGAVSSVASTLNAEHTGVVHAAVLGDNYPSKWKKGSGIDSWNMYVRQCTSFVAFRLSSANGFQLPKGYGNACTWRHIAKKQGYTVNKTPKVGAVAWFDTNAFQSHATYGHVAWVAEVRGDSVVIEEYNYNAGQGPEKYHKRQIPKNHVSGYIHFKDLPSGEASKSQTKEQQVSKEAVKQGGTYHFTERTPVKAQAQLTSPDLAYYNPGQSVHYDQAMTVDGHEWISYLSFSGNRRYIPIKKIEQKTQQVSETTSPINIGDRVTFPGVFRVDRIVNNLLVSEELAGGGATSLNWIDPSPLDETDRKGVKAGNQILQAGEFFVIPGNYRVLKVDRPSNGIYVKIGSRGTWLTADKASKLP

>WP_247211969.1 SH3 domain-containing protein [Streptococcus uberis]

MKFIKILLSQIVSLFLLLTISLHALETVNAAVLGDDYPIAWKSGWGTDTWGMYRRQCTSFVAFRLNQVNGFSIPSGLGNADTWGHIARRMGYPVNDIPAVGAVAWFDKGINGSHLAYGHVSWVAEVNGNLVTLEEYNFDAGQGPEQYHRRVINSHQVSGFIHFKDIDTGNIPISSPMQSPKSTIPNQGSYHFTEQMPIKAQPIANNQAIAFYQAGQMVHYDKTVIADGYQWLSYIAYSGQRRYIPIAKVTSKESSKQEDFAPGDQVTFSGVYQVTQIHGSLLSSKELAGGEPGPLNWLDPGPVLESNRDGQQSGDQILYPGDFFIIPGNYKVLQIHKETNGLLIQIGNRQTWVSMNKVQKST

>VEH28925.1 amidase [Streptococcus equi subsp. equi]

MLASGAILLGGNGAISSATTTISQGQSGIVHAAVLGDNYPSKWKQGFGADPWNMYLRQCTSFVAFRLHSSNGFSLPRGYGNVESWGRIAKHQGYHVDHTPRVGAVAWWDKGFNQSHALYGHVAWVAEVNGDTVIIEEYNYNAGQGPEKYHKRQIHKHQVSGYIHFKDLDHNSTSHTQMLHASQAGSTRLAKSGTYYFSSQSPIKAEATLASPDLAYYHTGQLVHYDQTLVADGYEWLSYIGHSGNHRYIPIHKLPVQPQQHSQASIDKPKPVATTPIKVGDTVAFPGVFRVDQVAQNMIASSELAGGEPTSLNWIDSAPLHETDHKGRIAGNQILQVGDYFVVSGTYKVLKVDQPSKGIYVQMGSRGTWLTIDKAKKV

>WP_206158765.1 M23 family metallopeptidase [Streptococcus equi subsp. zooepidermicus]

MKRIFFAFLSLCLFIFGTQTVSAATYTRPLDTGNITTGFNGYPGHVGVDYAVPVGTPVRAVANGTVKFAGNGANHPWMLWMAGNCVLIQHADGMHTGYAHLSKISVSTDSTVKQGQIIGYTGATGQVTGPHLHFEILPANPNWQNGFSGRIDPTGYIANAPVFNGTTPTEPTTPTTNLKIYKVDDLQKINGIWQVRNNILVPTDFTWVDNGIAADDVIEVTSNGTRTSDQVLQKGGYFVINPNNVKSVGTPMKGSGGLSWAQVNFTTGGNVWLNTTSKDNLLYGK

>WP_010921776.1 SH3 domain-containing protein [Streptococcus pyogenes]

MKKFHRFLVSGVILLGFNGLVPTMPSTLISQQENLVHAAVLGDNYPSKWKKGNGIDSWNMYIRQCTSFAAFRLSSANGFQLPKGYGNACTWGHIAKNQGYPVNKTPSIGAIAWFDKNAYQSNAAYGHVAWVADIRGDTVTIEEYNYNAGQGPERYHKRQIPKSQVSGYIHFKDLSSQTSHSYPRQLKHISQASFDPSGTYHFTTRLPVKGQTSIDSPDLAYYEAGQSVYYDKVVTAGGYTWLSYLSFSGNRRYIPIKEPAQSVVQNDNTKPSIKVGDTVTFPGVFRVDQLVNNLIVNKELAGGDPTPLNWIDPTPLDETDNQGKVLGDQILRVGEYFIVTGSYKVLKIDQPSNGIYVQIGSRGTWVNADKANKL

>WP_003083717.1 SH3 domain-containing protein [Streptococcus porcinus]

MKYFTNKGISFLLFVSLGLATLFPSCCQPVTAAVIGDDYPINWKSGLGSDTWGMYLRQCTSFVAYRLHIVNGFSLPSGYGNADSWGHVARRNGYQVDINPQVGSVAWFDKGVSLSHREYGHVAWVAEVKGDLVTLEEYNYNAGQGPEKYHRRQILRQEASGYIHFKNLINNSSNISISQLSRHSYLAPKGSYTFNQRKAVKAQPSLNSSDLVYYEKGNTVHYDKILDADGYRWLSYIGGSGNRRYIAIEKLEHQVETNDFKVEQIVHFKGPFQVSHNHGWMILSNILAGGAATQLNWLDPGPLIETDKIGQKAGDQVLYPGEYFSIPGEFKILQVDQESKGIKIKIGQKQTWISMDQVHKK

>WP_077323314.1 SH3 domain-containing protein [Streptococcus pseudoporcinus]

MKYFTNKGILFLLLISLGLTTLFPSCCQPVRARVIGDDYPINWKSGRGADSWGMYLRQCTSFVAYRLHNVNGFSLPPGYGNADSWGHVARRKGYQVDINPQVGSVAWFDKGVNLSHRDYGHVAWVAEVKGNLVTLEEYNYNAGQGPEYHRREILRQEASGYIHFKDLINNSSNTSISQLSRQLKLPSKGSYTFSQRRAVKAHPSLNSSDLIYYEKGNIVHYDKVLDVDGYRWLSYIGGSGNRRYVAIEKLEQQVSPREFQVEQIVQFKGSFQVSHNHGWMISSDILAGGPATQLNWLDPGPLIETSKNGQKAGDQVLYPGDYFSIPGEFKILQIDQKSKGIKIKIGQKETWVSMDHVHQKAS

>WP_008089638.1 SH3 domain-containing protein [Streptococcus ictaluri]

MKKFHQLLVSGAILLSVNGAISTAAPLFNPNSVSQVQAEVLGDNYPSKWKKESGIDSWNMYVRQCTSFVAFRLSSANGFQLPRGYGNAHTWGHIAKTQGYAVNKTPKVGSVAWFDKGANNSSAIYGHVSWVAEVSNDYVTLEEYNYNAGQGPEKYHKRQIHKSKVSGFIHFKDLPSTTSSSSVEINQPLSNKDKTLPKSGLYQFTQKHLIKGEAKIDSPDLGYYDAGQSVNYDQLLSSDGYDWLSYLSFSGNRRYIPLKKITEKSNTVEVTHAKTTTTTAIQNPIVVGDQVTFPGVFKVTQVINGLITSSQLAGGDPTHYNWIDPGPLTETDAKGNTIGDQILHSGEYFTVPGTYKVLKVDKPSNGIYVQMGSRGTWFIMDKAKKV

>WP_003045998.1 SH3 domain-containing protein [Streptococcus canis]

MKKCHQFLVSGAILLSVNGAVSAVATALKPEQTGIVHAAVIGDNYPSKWKKGTGIDSWNMYVRQCTSFVAFRLSSANGFQLPRGYGHAYTWGHIAKKQGYSVNKTPKVGAVAWFDTDAYQSNPIYGHVAWVAEVRGDQVIIEEYNYNAGQGPEKYHKRQIPKHQVSGYIHFKDLPADANHSQEKQQPATQTPTNQSGTYHFTTQVPIKAQAQLASPDLAHYSAGQSVHYDQILTAEGYEWLSYLSFAGNRRYIPIKKLSQSTSQVPETKLPINVGDTVTFPGVFRVDRIVNNLLVNEELAGGDATSLNWLDPTPLDETDNNGKVSGDQIFRIGESFIVPGHYKVLKVDRPSNGIYVKIGSRGTWLSADKANKV

>WP_099567791.1 SH3 domain-containing protein [Streptococcus parauberis]

MKKFQTLIVLAVFVSQLVNPFTLNTKAAVLGDNYPSQWQTGWGPDTWGMYKRQCTSFVAFRLSSANGFTLPTGYRNADSWGHIARSQGYSVDMQATVGAVAWFDKGVNYSHQDYGHVAWVAEVNGNQVTLEEYNYNAGQGPEKYHRRTIAMGAVSGYIHFKDLGQSHSNPTPTAAPANLAPSGTYYFTERTAVRPEAKLNSHALTHYEKGQSVIYDRVLTAEGYQWLSYIGGSGNRRYVAIKKLASSVKPKDPETKPSQFKVGDQVTFPGVFKVNQHHGSLISSVDLAGGSPTSLNWIDPTPVDESNQQGQKAGDQFLNPGEYFIIPGKFKILQIDQKTNGIRVQIGSRATWVTMARAKKA

>WP_054278001.1 SH3 domain-containing protein [Streptococcus phocae]

MKTYRQFLVSGAILLSLNGAVSSLASPFTTTPSGVVHAAVLGDNYPTKWKKGFGIDSWNMYLRQCTSFVAFRLSSANGFSLPRGYGNADSWGHIARNQGYQVNKTPKIGAVAWWDKGINQSHALYGHVAWVADIKGDYVTIEEYNYNAGQGPEKYHKRTIHKNHVSGYIHFKDLNTIVSHQVQPISQEKPETATSLAQSGTYHFTSKSPIKGEPKLSSPDLTHYDTGQSVHYDQVVTAEGYEWISYLSFSGNRRYIPIRKIAEQPSPSKPSVSETVTVNITVGDTVSFPEAFQVTSVQGNLVTSASLAGGDPTHYNLIDPTPVNETDAAGNITGDQILRPGEYFSIPGTYKVLKVDKPSNGIYVQIGSRGTWVSTDKAKKI

>WP_071127360.1 SH3 domain-containing protein [Streptococcus iniae]

MKILKQVLSLTLVLTSLWGIQTLPLTNKSLAFSKTVKAAVLGDNYPHKWQKGWGADSWGMYLRQCTSFVAFRLSTANGFNLPAGYGNADSWGHIAKAQGFLVNQIPQVGAVAWFDKGVNYSHRDYGHVAWVAEVSGDYVTLEEYNYNAGQGPEKYHRRQIHRQQVSGFIHFKDITTEQSTIPSSKHILAESTKLLPNKGTYYFKAMSPVKEEARFQSLTLTSYSAGQYVHYDQTMVADGYQWISYIGFSGNRRYIPVVKIEERKSESPETPKSERIFKVGDTVSFQGTFKVTQIHGNLVSSADLAGGQPSQLNWIDPGPVDKTNASGQKEANQILSPDNYFTIPGTYKVLQVDLPSRGLYIQIGSRASWVTMDRVLKQ

>WP_011225268.1 CHAP domain-containing protein [Streptococcus thermophilus]

MKKKLLNYLILLACLATVSVSCSGSAQAAVLGDDYPFSWKYGGFGVDPWTMYWRQCTSFAAYRLSNTNGFTLPIGYGNAITWGNIARANGHRVDMNPAVGSIAWFSAGVNGAGHMGHVAWVAEVHGDQVTIEEYNYDAGQGPEKYHKRSFHKSQVSGYIHFKDLESGTQNGNPTNSSIKVGDSVRFTGTFRVTSVSGNTITSQDLAGGTPTKHNIVDPSPVLEVDAQGNPTSDQYLNPGETFTIPGNFKVLAIDPPSDGILVQIGNLKTWVTQSVLEKI

>WP_175060955.1 CHAP domain-containing protein [Streptococcus salivarius]

MKKKLLNCLLPLACLATVSVSCGSSAQAAVLGDDYPSSWKYGGFGVDPWTMYWRQCTSFAAYRLSNTNGFTLPVGYGNAITWGSIARANGHRVDMNPAVGSIAWFSAGVNGAGHMGHVAWVAEVHGDQVTIEEYNYDAGQGPEKYHKRSFHKSQVSGYIHFKDLEPGAQNGNSTNSSIKVGDTVRFTGTFRVTSVSGNTITSQDLAGGTPTKHNIVDPGPVLEVDGQGNPTSDQYLNPGETFTIPGNFKVLAIDPPSDGILVQIGNRKTWVTQSVLEKV

>WP_003094950.1 CHAP domain-containing protein [Streptococcus vestibularis]

MKKKLLNCLLPLACLATISVSCGSSAQAAVLGDDYPSSWKYGGIGVDPWTMYWRQCTSFAAYRLSNSNGFTLPVGYGNAITWGPIARANGHRVDMNPAVGSIAWFSNGVNGAGYMGHVAWVAEVNGDQVTIEEYNYNAGQGPEKYHKRSFHKSQVSGYIHFKDLEPGAQNGNSTNSSIKVGDTVRFSGTFRVTSVSGNTITSQDLAGGTPTKHNIIDPGPVLEVDGQGNPTSDQYLNPGETFTIPGNFKVLAIDPPSDGVLVQIGNRKTWVTQSVLEKV

>WP_012775400.1 SH3 domain-containing protein [Streptococcus suis]

MKTFLRRKGIACQLLAAIFIVGINLQSPPSASANARGDDYPYTAVDAVDPWRLYTRQCTSFVAFRLSTVNGFTLPPAYGDANVWGHRAKQEGYQVNMTPAVGAVAWWETPMHVAWVSAVNGDTVEIEEYNYGVRYTYGRRVIPKNAVSGYIHFKNLGAGTTNSQPALAPSGTYTFTSQLAIKSQASLSSPTLDYYYAGESVRYDKVLEDDGQRWISYISYSGARRYIAIQPTITQSTNNSLPASGSYTIQSYSSVRNNPSLSSPEITFYGAGSVVRYDKVITAEGRTWISYISYSGARRYIAIS
